# Yttrium-90-doped metal-organic frameworks (MOFs) for low-dose rate intratumoral radiotherapy

**DOI:** 10.1101/2024.09.14.613012

**Authors:** Xiaoli Qi, Anzhelika Fedotova, Zhihao Yu, Anna Polyanskaya, Ningfei Shen, Bayirta Egorova, Dmitry Bagrov, Tatiana Slastnikova, Andrey Rosenkranz, Gilles Patriarche, Yurii Nevolin, Anastasia Permyakova, Stanislav Fedotov, Mathilde Lepoitevin, Stepan Kalmykov, Christian Serre, Mikhail Durymanov

**Author notes:** Corresponding author*, **Dr. Mikhail Durymanov** – Medical Informatics Laboratory, Yaroslav-the-Wise Novgorod State University, Ul. Bolshaya St. Peterburgskaya, 41, Veliky Novgorod, 173003, Russian Federation. Equal contribution (Xiaoli Qi, Anzhelika Fedotova, Zhihao Yu).

## Abstract

Brachytherapy, or intratumoral radiation therapy, is a highly effective treatment option for localized tumors. Herein, we engineered injectable and biodegradable metal-organic frameworks (MOFs) to deliver the therapeutic radioisotope yttrium-90 (^90^Y). Particles of bimetallic MIL-100(Fe,Y) and Y-BTC, doped with ^90^Y and ^88^Y, were synthesized in a single step and retained radioyttrium in various buffer solutions. Tumor injectability and radioisotope retention were evaluated using tumor-bearing mice. *In vivo* analysis and calculations showed that radiolabeled MIL-100(Fe,Y) emitted more than 38% of its radioactivity, while Y-BTC emitted greater than 75% of its radioactivity, through 7 days at the tumor site upon intratumoral injection, without significant yttrium accumulation in off-target tissues. The anticancer effects of MIL-100(Fe,Y,^90^Y) and ^90^Y,Y-BTC particles were assessed using 3D multicellular tumor spheroids and a tumor-bearing mouse model, respectively. ^90^Y-doped MIL-100(Fe,Y) particles penetrated A549 tumor spheroids and caused superior cytotoxic effects compared to non-radioactive particles or ^90^YCl_3_, added at the same dose. Brachytherapy with ^90^Y-doped Y-BTC MOFs induced inhibition of B16F1 melanoma tumor growth and resulted in an increased median survival of 8.5 days compared to 4.5 days in untreated mice. This study exhibits the feasibility to prepare radioactive ^90^Y-containing biodegradable, non-toxic MOF particles that are advantageous for low-dose rate internal radiotherapy.

## Introduction

Low-dose rate (LDR) internal radiation therapy, commonly known as brachytherapy, stands as a highly effective treatment modality for localized tumors^[1]^ and presents a promising strategy for addressing medically inoperable malignancies.^[2]^ This therapeutic approach implies the local delivery of radionuclide carriers directly into the tumor, allowing for the administration of maximum effective radiation dose while minimizing damage to surrounding healthy tissues.

However, conventional radioactive implants, such as tiny titanium ‘seeds’ with a length of several millimeters, exhibit several drawbacks. Owing to their inert nature, they permanently retain in the body and may lead to pain and discomfort for patients, including urinary incontinence and erectile disfunction.^[3]^ Additionally, issues such as bleeding and inflammation upon implantation, as well as the potential for migration into circulation with a risk of embolism,^[4,5]^ further underscore the need for improved alternatives.

To address these challenges, several biodegradable radionuclide carriers of different nature have been developed. For example, cross-linked chitosan implants encapsulated with ^131^I-norcholesterol have been designed as potential degradable devices for brachytherapy.^[6]^ Other reported polymer-based implants include thermogelling co-polymer systems with encapsulated yttrium-90^[7]^ and polypeptide cross-linked hydrogels with ^131^I-labeled tyrosine residues.^[8]^ In addition, some types of biodegradable glasses were developed for fabrication of biodegradable seeds.^[9]^ All the mentioned examples emphasize the use of radioisotopes with short half-lives and materials that degrade slowly, with the aim of maximizing the emission of high radiation doses within tumor tissue while mitigating the risk of radionuclide leakage and ensuring the implant’s biodegradability. Despite advancements, injectable biodegradable implants face significant concerns, such as poor retention and off-target toxicity due to premature degradation.^[8]^ Additionally, the lack of diversity in materials and designs limits their ability to meet the personalized treatment needs of different tumors.

Metal-organic frameworks (MOFs), a versatile class of ordered porous hybrid solids composed of metal sub-units and organic linkers, emerge as ideal candidates for radionuclide incorporation. In contrast to conventional inorganic particles, MOFs are intrinsically biodegradable due to the adjustable coordination interactions between metal sites and organic linkers, and some MOF structures exhibit slow-rate biodegradation behavior in cells^[10]^ and *in vivo*.^[11]^ These materials, once the appropriate composition is selected, can be poorly toxic and biodegradable^[12,13]^ while their synthesis can be achieved under green and scalable conditions.^[14]^ Moreover, tunable MOF porosity can be exploited for the encapsulation of anti-cancer drugs^[15,16]^ for combined efficacy that adds another layer of advantage to such materials. Previous studies have evaluated the retention of radionuclides, diagnostic potential, and therapeutic effects of certain MOF structures doped with radionuclides such as ^89^Zr,^[17]^ ^99m^Tc,^[18]^ and ^177^Lu.^[19]^

Due to the high-energy β-particles emitted by ^90^Y, which have a maximum energy of 2.28 MeV and can penetrate deeply into tissues, this radionuclide is widely used for both cancer and non-cancer applications. Another radioisotope of yttrium, ^88^Y, has a longer half-life (106.6 days compared to 64.1 h for ^90^Y), allowing for extended periods for radionuclide biodistribution measurements. To our knowledge, the integration of radioyttrium into MOFs has not yet been reported. One can imagine that ^88^Y and ^90^Y could be seamlessly incorporated into Y-based MOFs, such as the Y-BTC structure (BTC for Benzene-1,3,5-TriCarboxylate), that is produced through a rapid water ambient temperature protocol, highly suitable here to load radioactive ^90^Y inside the MOF particles by direct synthesis and avoid any toxicity effects.^[20,21]^ Such as rapid synthesis is required as one aims here to load radioactive Y inside the MOF without losing too much of the radioactivity thus requiring short synthesis times, ideally less than 12-24 h. Non-yttrium-based biocompatible MOF structures, such as mesoporous Fe-BTC MOF nanocarrier MIL-100(Fe), can also be synthesized at low pressure in water, within less than one day, without any additives or toxic products^[22]^ and seem promising for the coordination of yttrium and the chemistry of Y^3+^ is to some extent compatible, despite a slightly larger ionic radius,^[23]^ with the one of Fe^3+^, authorizing the partial substitution of Fe ions by Y cations inside MIL-100 through a direct synthesis route. Moreover, this MOF addresses the limitation of Y-BTC’s pore channels, that are too narrow to load drugs.

In this study, we have therefore evaluated the use of carboxylate-based MOFs as slowly biodegradable, biocompatible, and injectable carriers for intratumoral delivery of beta-emitter yttrium-90 (^90^Y), which is commonly used in brachytherapy for treating various types of cancers. We obtained two ^88^Y- and ^90^Y-doped MOF trimesate-based structures using a one-pot reaction in water under ambient pressure. First, yttrium radioisotopes were incorporated into Y-based microporous Y-BTC MOF,^[24]^ resulting in rod-like particles. In the case of the mesoporous iron-based MIL-100 MOF,^[14]^ the yttrium doping resulted in bimetallic MIL-100(Fe,Y) particles. Both types of MOFs contained less than 10^-3^(for MIL-100)-10^-5^(for Y-BTC) % of radioyttrium relative to the maximum capacity for yttrium, allowing for the adjustment of specific activity across a broad range to match the desired injection volume and MOF concentration. We conducted a comprehensive characterization and investigation of the physicochemical properties of both Y-doped MOFs. In particular, we employed Differential Pair Distribution Function (dPDF) analysis to elucidate the structural modifications in MIL-100(Fe) induced by Y-doping. For both types of ^88^Y-doped MOFs, we then evaluated radioisotope retention in tumor tissue upon local injection demonstrating low rate of radioyttrium washout from the tumor. Finally, the therapeutic effects of MIL-100(Fe,Y,^90^Y) and ^90^Y,Y-BTC particles were estimated using 3D multicellular tumor spheroids and tumor-bearing mouse model, respectively. Both types of radioactive particles exhibited anticancer effects, demonstrating potential for use in low-dose rate internal radiotherapy.

## Materials and methods

### Synthesis of ^90^Y-doped MOFs

MIL-100(Fe,Y) and Y-BTC were synthesized following the previously reported procedures but substituting part of the Fe salt by Y salt.^[23,25]^ For the preparation of MIL-100(Fe,Y), 0.57 g of H_3_BTC (Sigma Aldrich) and 0.32 g of NaOH (Sigma Aldrich) were added to 50 mL of deionized water, and the solution was sonicated until all the H_3_BTC dissolved. In the second solution, 0.81 g of FeSO_4_·7H_2_O (Sigma Aldrich) and an appropriate amount of Y(NO_3_)_3_·6H_2_O (Acros Organics) were dissolved in 50 mL of deionized water, achieving n(Y)/[n(Y)+n(Fe)] ratios of 0%, 2%, 5%, 8%, 10%, and 15% respectively. Both solutions were then mixed and allowed to react in the presence of air for 24 h at 60°C. The resulting precipitate was separated from the supernatant liquid by centrifugation at 5,600×g for 10 min and washed with 3×50 mL of deionized water and 3×50 mL of denatured ethanol. The product was finally dried in an oven at 40°C. For the preparation of MIL-100(Fe,Y,^88/90^Y), either ^88^YCl_3_ (Ritverc Gmbh) or ^90^YCl_3_ was used. The isolation of ^90^Y was carried out from a ^90^Sr solution using chromatographic Sr-resin (Triskem inc.). MIL-100(Fe,Y,^88/90^Y) particles were synthesized according to the same protocol, with the addition of either 200 μL of ^88^YCl_3_ with specific activity of 113 kBq mL^-1^ or 500 μL of ^90^YCl_3_ with specific activity of 16.6 MBq mL^-1^ to the second solution. The radioactivity of ^88^Y was measured by a gamma spectrometer with a HPGe-detector GR3818 Canberra Ind. The radioactivity of ^90^Y was measured by a liquid scintillation counter Perkin Elmer Tri-Carb 2810 TR.

For the preparation of Y-BTC, 1.2 g of H_3_BTC and 0.6 g of NaOH were added to 30 mL of a water–ethanol mixture (v/v = 1:2), and then sonicated until all the H_3_BTC dissolved. Next, 1.9 g of Y(NO_3_)_3_·6H_2_O was dissolved in 20 mL of a water–ethanol mixture (v/v = 1:2). The H_3_BTC solution was added to the Y(NO_3_)_3_·6H_2_O solution at room temperature. The mixture was further vigorously stirred for 24 h. The product was collected by centrifugation at 5,600×g for 10 min, washed three times with water and ethanol alternately. Finally, the product was dried in the oven at 40°C. Particles of ^88/90^Y,Y-BTC were synthesized similarly, with the addition of either 200 μL of ^88^YCl_3_ with a specific activity of 113 kBq mL^-1^ or 500 μL of ^90^YCl_3_ with specific activity of 16.25 MBq mL^-1^ to the Y(NO_3_)_3_·6H_2_O solution.

### Characterization of ^90^Y-doped MOFs

The phase composition of the Y-BTC samples was determined based on powder X-ray diffraction (PXRD) data collected on a Huber Guinier camera G670 equipped with an Image Plate detector. Co-K_α1_ radiation (λ = 1.7899 Å) and a curved Ge(111) monochromator were used, covering a 2θ range from 4 to 100°. The simulated pattern was obtained from the Cambridge Crystallographic Data Centre (CCDC). The unit cell parameters were refined using Jana2006 software.^[26]^

PXRD data for MIL-100(Fe,Y) were recorded using a high-throughput Bruker D8 Advance diffractometer working in a transmission mode and equipped with a focusing Göbel mirror producing Co-K_α1_ radiation (λ = 1.5418 Å) and a LynxEye detector.

The solid solutions of MIL-100(Fe,Y) (with 0 to 10 % of Y) were studied by PXRD using Mo-K_α_ radiation (λ = 0.71073 Å). The powders were packed to the 0.7 mm glass capillaries and measured using Empyrean diffractometer (Malvern Panalytical) with GaliPIX^3D^ hybrid pixel detector and measured from 2 to 145 2θ°. The unit cell parameters of the obtained diffraction patterns were fitted by the Le Bail method in Fullprof.^[27]^ The differential pair distribution function analysis (dPDF) was performed using PDFGetX3 software.^[28]^ For that purpose the PDFs of the Y-doped samples were normalized with MIL-100(Fe) on trimesate C-C (∼2.60 Å) correlation. From the normalized PDFs of MIL-100(Fe,Y) the non-doped MIL-100(Fe) PDF was subtracted in order to see the appearance of novel interatomic correlations (Y-O bonds).

The biodegradability of Y-BTC and MIL-100(Fe,Y) was tested after dispersing them in Phosphate-Buffered Saline (PBS, pH 7.4) and Fetal Bovine Serum (FBS) at 37°C. The concentration of samples is 5 mg mL^-1^. At different times (1, 2, 4, 7, 10 days), the suspension of particles was centrifuged (13,500 rpm, 10 min) to collect the solid, followed by washing with Milli-Q® water. Then the crystallinity of these solids was analyzed by PXRD.

Thermogravimetric analysis combined with differential scanning calorimetry (TGA/DSC) measurements under oxygen were recorded on a Netzsch STA Jupiter 449F3 system (Germany). Gas flows of 100 mL min^-1^ and heating rates of 3 K min^-1^ were employed.

Scanning electron microscopy coupled with energy-dispersive X-ray spectroscopy (SEM-EDX) results were recorded with an FEI Magellan 400 scanning electron microscope. Fe and Y contents were averaged over 5 points for each sample. Scanning Transmission Electron Microscopy - High Angle Annular Dark Field (STEM-HAADF) images were obtained from a Titan Themis 200 microscope operating at 200 kV. The microscope was equipped with a Ceta 16M hybrid camera capable of working under low electron irradiation conditions. The images were acquired in low-dose conditions with an irradiation current between 100 and 250 electrons per square angstrom. For the TEM grid preparation, a 10 μL drop of the sample solution was placed on a 200-mesh copper grid covered with a pure carbon membrane (from Ted Pella).

FT-IR spectra were measured with Nicolet iS5 FTIR ThermoFisher spectrometer in the 3,500-600 cm^-1^ range.

Particle size measurement of hydrodynamic diameters and charge analysis were performed on a Malvern Zetasizer NanoZS (Malvern Instruments). Samples for measurements were prepared at a final concentration of 0.1 mg mL^-1^.

N_2_ sorption isotherms were obtained at 77K using Micromeritics TriStar II PLUS. Prior to the analysis, approximately 50 mg of the samples were activated for 8 h at 150°C under primary vacuum. Brunauer-Emmett-Teller (BET) surface area were estimated at a relative pressure lower than 0.25.

### In vitro release of radioyttrium from MIL-100(Fe, Y,^88^Y) and ^88^Y,Y-BTC particles

To measure the *in vitro* radioyttrium release profiles of Y-doped MOFs, MIL-100(Fe,Y,^88^Y) and ^88^Y,Y-BTC particles were dispersed in different physiologically relevant liquids including Milli-Q® (deionized) water, 0.1 M (pH 7.5) 4-(2-hydroxyethyl)-1-piperazineethanesulfonic acid (HEPES), PBS (pH 7.4), and FBS (Gibco). Briefly, 60 mg of Y-doped MOF particles was dispersed in 10.5 mL of each solution and kept in the incubator at 37°C with mild shaking. At different incubation time points, 5 mL of supernatant was collected after centrifugation of the suspension for 10 min at 17,000×g, and replaced with the same volume of fresh buffer solution. The radioactivity of the yttrium was measured using a gamma spectrometer.

### Fluorescent labeling of MIL-100(Fe,Y) particles

Prior to conducting penetration and uptake investigations in 3D spheroids, the MIL-100(Fe,Y) particles were fluorescently labeled with toluidine blue O (TBO) dye using the sulfo-NHS/EDC reaction. First, 25 mg of MIL-100(Fe,Y) particles were dispersed in 6 mL of 5 mM HEPES buffer (pH 7.4) and sonicated for 2 min. Then, 8 mg of 1-ethyl-3-(3-dimethylaminopropyl)carbodiimide hydrochloride (EDC) and 13.2 mg of N-hydroxysuccinimide (NHS) were dissolved in 3.5 mL of HEPES buffer and added to the MIL-100(Fe,Y) particle suspension, followed by stirring for 30 min at room temperature. Next, 500 μL of TBO solution (0.5 mg mL^-1^) in anhydrous DMSO was added to the mixture and stirred for 2 h at room temperature. After the reaction, the TBO-labeled particles were collected by centrifugation at 20,000×g for 20 min and washed 3-4 times with ethanol and water to remove unreacted TBO. Finally, the TBO-labeled MIL-100(Fe,Y) particles were lyophilized for 24 h.

### Generation of tumor spheroids

Multicellular tumor spheroids were generated by two methods. For production of A549 spheroids, we used 9×9-well MicroTissues 3D Petri Dish® micro-molds as described earlier.^[29]^ Briefly, the micro-molds were sterilized with anhydrous ethanol and allowed to dry under UV light for 30 min. Then, they were filled with sterile 2% (w/v) agarose solution, prepared in Milli-Q® water. After solidification, the gelled agarose molds were released from flexible 3D Petri Dish® micro-molds and transferred to 6-well plates (3 agarose molds per well). To equilibrate the agarose gels, each well was filled with 1 mL of growth medium. After equilibration, the plates were placed into the incubator and incubated overnight. Before seeding the cells, the medium was removed from both the culture plate and the molds. Then, 1.3 mL of fresh growth medium was added to each well and aliquots of 150 μL of growth medium with 40,000 cells were gently added into the molds. The medium was changed every other day.

To generate B16F1 spheroids, the cells were plated in round-bottom 384-well plates with an ultra-low attachment coating (Corning, Kennebunk, ME) at a density of 1,000 cells per well in 25 μL of the medium. The spheroids were incubated at 37°C under humidified atmosphere with 5% CO_2_.

The size and morphology of the spheroids were examined using bright-field light microscopy with an AxioVert.A1 microscope (Zeiss, Oberkochen, Germany) equipped with a Plan 10×/0.25 phase-contrast objective.

### MIL-100(Fe,Y) particle uptake by tumor spheroids

To study the uptake and penetrating ability of MIL-100(Fe,Y) particles, we used A549 spheroids. On day 4, the spheroids were treated with TBO-labeled MIL-100(Fe,Y) particles at a final concentration of 100 μg mL^-1^. Following 12- and 24-h incubation periods, the 3D spheroids were harvested, embedded in HistoPrep 355 tissue embedding media (Fisher Scientific, Ottawa, Canada), and frozen at −20°C. The frozen blocks were then cut into 10-μm sections, fixed in a 1:1 mixture of acetone and methanol for 15 min, and air-dried at room temperature. Nuclear DNA was stained with DAPI for 10 min. Images of the spheroid sections were acquired in multitrack mode using a Zeiss LSM 710 confocal microscope (Carl Zeiss, Oberkochen, Germany) equipped with a Plan-Apochromat 63×/1.4 Oil DIC lens.

### Cytotoxicity analysis of MIL-100(Fe,Y,^90^Y) particles using A549 spheroids

The cytotoxicity of MIL-100(Fe,Y,^90^Y) particles was investigated using the A549 lung spheroid model. After 4 days of incubation at 37°C in a humidified 5% CO_2_ atmosphere, the spheroids were harvested and distributed among the wells of a non-attachment round-bottom 96- well plate (10 spheroids per well). On the same day, MIL-100(Fe,Y,^90^Y) particles were added to the first group of wells with spheroids at concentrations of 30 µg mL^-1^ (0.12 µCi per well), 75 µg mL^-1^ (0.3 µCi per well), and 100 µg mL^-1^ (0.4 µCi per well). Non-radioactive MIL-100(Fe,Y) particles were added to the second group of wells with spheroids at concentrations of 30 µg mL^-1^, 75 µg mL^-1^, and 100 µg mL^-1^. Finally, a ^90^YCl_3_ solution was added to the wells of the third group containing spheroids at doses of 0.12 µCi per well, 0.3 µCi per well, and 0.4 µCi per well (less than 10^-10^ M YCl_3_). On day 5 after the addition of the formulations, the viability of the treated and non-treated A549 spheroids was assessed using the CellTiter-Glo® 3D Cell Viability Assay (Promega, Madison, WI, USA) according to the manufacturers’ instructions.

### Cytotoxicity of ^90^Y measured in B16F1 spheroids

The cytotoxicity of ^90^YCl_3_ was investigated using the B16F1 3D spheroid model. The day after seeding, the spheroids were treated with 25 µL of the growth medium containing ^90^YCl_3_ at concentrations of 20 µCi mL^-1^ (approximately 19.9 Gy per well), 10 µCi mL^-1^, 5 µCi mL^-1^, 1 µCi mL^-1^, and 0 µCi mL^-1^ (5 wells for each dose). Molar concentration of YCl_3_ was less than 10^-9^ M. The spheroids were photographed daily, and the growth medium (25 µL) was changed every other day starting 72 h post-treatment.

### Animal models

C57BL/6 mice (8-week-old females) were used for all animal experiments in this study. The mice had access to food and water *ad libitum* and were housed at room temperature. The work with animals was performed in accordance with the EU Directive 2010/63/EU for animal experiments and was approved by the Bioethics Commission of Lomonosov Moscow State University, meeting no. 152-d held on 18.05.2023, application no. 151-a. The tumor-bearing mouse model was established by subcutaneous injection of 1×10^6^ of LLC-1 cells or 1×10^6^ of B16F1 cells in a volume of 50 µL of HBSS into the right flank for each mouse, using a 29G1/2 needle and a 1-mL syringe. Tumor growth was measured daily using a caliper, and the tumor volume was calculated using the formula: volume = (tumor length) × (tumor width)^2/2.

### Intratumoral retention and biodistribution study

Tumor retention of ^88^Y was studied at different times after intratumoral administration of MIL-100(Fe,Y,^88^Y) and ^88^Y,Y-BTC MOF particles in C57BL/6 mice with LLC-1 and B16F1 tumors, respectively. Ten days after LLC-1 tumor establishment, each tumor-bearing mouse was anesthetized with xylazine and then received a single dose of MIL-100(Fe,Y,^88^Y) (1.25 kBq or 0.034 μCi in 40 μL) injected into tumors with volume of 400-900 mm^3^, using a 29G1/2 needle and a 1-mL syringe. Three groups of mice (3 animals each) were sacrificed at different time points (1, 3, and 7 days) after injection.

Similarly, ten days after tumor establishment, B16F1 tumor-bearing mice were treated with ^88^Y,Y-BTC particles (1.8 kBq or 0.049 μCi in 40 μL) injected into tumors measuring 100-300 mm^3^ using a 1-mL syringe after anesthetization with Zoletil®-xylazine combination. The mice (3 animals per time point) were sacrificed at 1, 3, 7, and 14 days post-injection.

Subsequently, tumors as well as organs including the liver, lungs, heart, kidneys, guts, bladder, and whole-body carcass were collected. The weight of each organ was determined and the level of radioactivity was quantified using a gamma spectrometer with HPGe-detector GR3818 Canberra Ind. Different organ radioactivity levels were calculated by determining the percentage of injected doses per gram of tissue (% ID per g) and per organ (% ID).

### In vivo therapeutic effects of ^90^Y,Y-BTC MOFs

Obtained Y-BTC and ^90^Y,Y-BTC (∼10.5 MBq or 284 µCi per 50 mg) particles were re-suspended in 0.9% NaCl solution to a concentration of 100 mg mL^-1^. C57BL/6J mice bearing B16F1 tumors were used for the experiment when the tumor volume reached approximately 200 mm^3^. The mice were randomly divided into 3 groups (8 mice per group). Group 1 received an intratumoral injection of the ^90^Y,Y-BTC particle suspension at a dose of 9.2 MBq (249 µCi) per gram of tumor. Group 2 received the same amount of the Y-BTC suspension. The control group (group 3) was injected with the same volume of saline. Post-injection, tumor volumes were measured daily and calculated using the formula: tumor volume = (tumor length)×(tumor width)^2/2. The animals were sacrificed when the tumor volume exceeded 1 mL. Tumors and organs, including the lungs, spleen, liver, bone marrow, and heart, were harvested from some animals for histological analysis.

### Statistical analysis

The experimental data analysis was carried out using GraphPad Prism 5 (GraphPad Software Inc., San Diego, CA). The data are presented as mean ± standard deviation (SD). Each experiment was performed at least in triplicate, and the number of samples or animals is additionally specified in the figure captions. To determine the statistical significance of differences between control and experimental groups, a one-way ANOVA followed by a post hoc Dunnett’s test was performed. Survival differences between animal groups were determined using a log-rank Mantel-Cox test. In all cases, a value of p < 0.05 was considered to indicate a statistically significant difference.

## Results

### Synthesis and characterization of Y-BTC MOF particles

Y-BTC (**Fig. 1A**) was synthesized using a previously reported eco-friendly method^[25]^ by mixing Y(NO_3_)_3_·6H_2_O, trimesic acid and NaOH in a water/ethanol mixture at room temperature for 24 h, followed by centrifugation and washing with water and ethanol.

**Figure 1.**
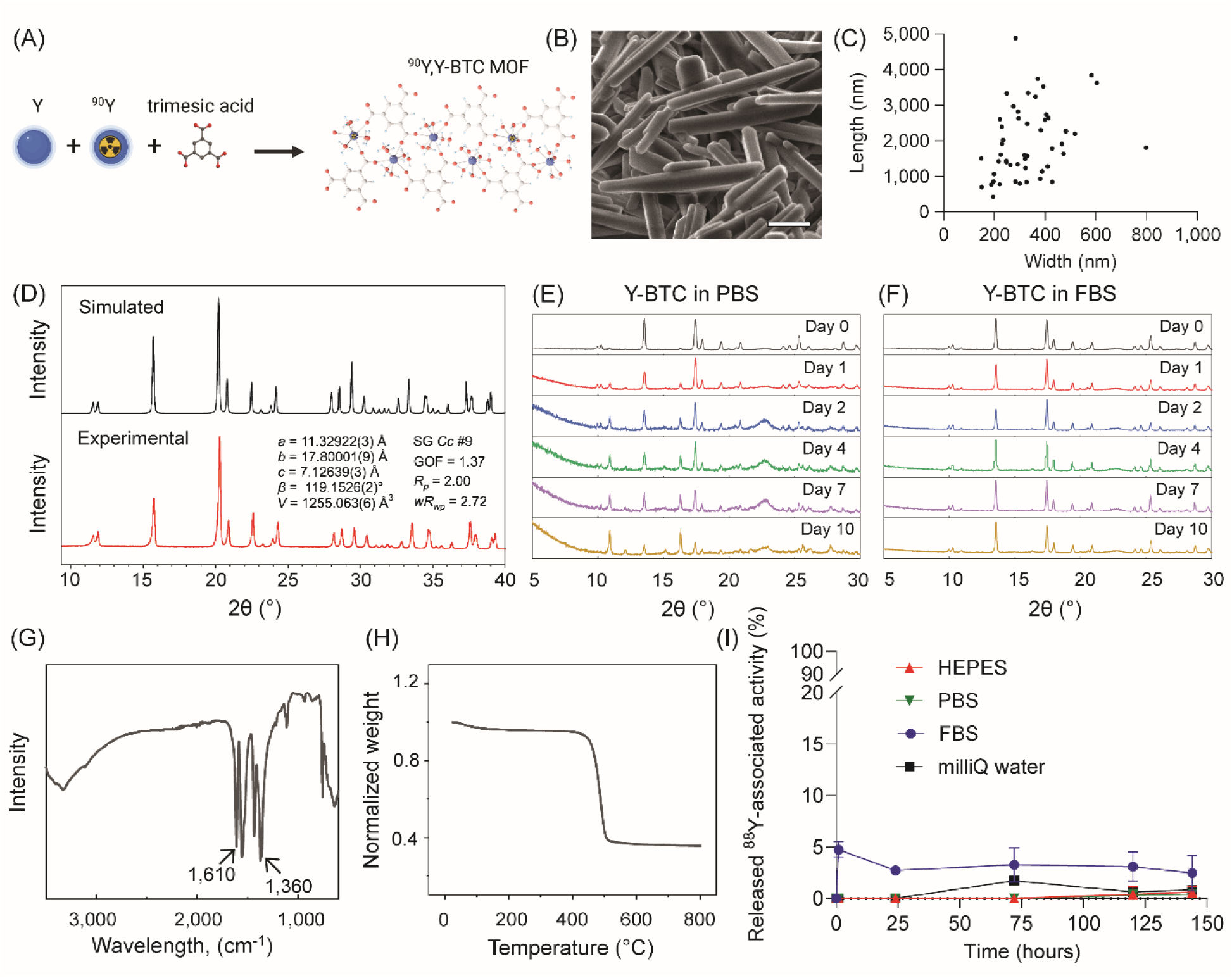
Physicochemical properties of Y-BTC MOF particles. (A) Scheme of Y(BTC)(H_2_O)_6_ MOF structure (denoted Y-BTC) after doping with radioyttrium. The denoted atoms are Yttrium (blue), Oxygen (red), and Hydrogen (light blue). (B) SEM images of Y-BTC particles. The scale bar is 500 nm. (C) Size distribution of Y-BTC particles determined by quantitative analysis of SEM images. Over 30 particles were measured. (D) PXRD pattern of simulated Y(BTC)(H_2_O)_6_ and experimental Y-BTC particles (Co-K_α1_ radiation). (E,F) PXRD patterns of Y-BTC soaked in PBS (E) and FBS (F) over 0-10 days at 37°C (Co-K_α1_ radiation). (G) FT-IR spectrum of Y-BTC. (H) Thermogravimetric curve of Y-BTC particles under oxygen flow (3°C/min heating rate) with the mass at 25°C normalized to “1”. (I) *In vitro* yttrium-88 release from ^88^Y,Y-BTC MOFs in different solutions, including HEPES (4-(2-hydroxyethyl)-1-piperazineethanesulfonic acid) buffer, PBS (phosphate-buffered saline), FBS (fetal bovine serum), and purified MilliQ® water, at different time points (1, 24, 72, 96, 120, and 144 h) of incubation at 37⁰C.

The Scanning Electron Microscope (SEM) image (**Fig. 1B**) showed an elongated rod shape with particle widths ranging from 140 to 800 nm and lengths ranging from 420 to 4,900 nm (**Fig. 1C**, **Fig. S1**).

The crystallinity of the Y-BTC or Y(BTC)(H_2_O)_6_ was studied by Powder X-ray Diffraction (PXRD) analysis. The PXRD patterns are in good agreement with the theoretical ones determined from single crystal structural data (cif) of Y-BTC using the Cambridge Crystallographic Data Centre (CCDC) Mercury software. Overall, the PXRD pattern (**Fig. 1D**) confirmed the single phase Y(BTC)(H_2_O)_6_ with refined cell parameters: S.G. *Cc*, *a* = 11.32922(3) Å, *b* = 17.80001(9) Å, *c* = 7.12639(3) Å, β = 119.1526(2)°, *V* = 1255.063(6) Å^3^ (**Fig. S2**), which are consistent with the literature data.^[20]^ Y-BTC slowly degraded in PBS (pH 7.4) revealing minor peaks at 10.8° and 16.3° (**Fig. 1E**), indicating the presence of an impurity phase corresponding to a MIL-78-type Y(BTC).^[30]^ Within 10 days of soaking in PBS, Y-BTC partially lost its crystallinity over time, whereas the one of MIL-78 was still observed, evidenced a better chemical stability for this MOF. However, Y(BTC)(H_2_O)_6_ demonstrated in FBS greater stability than in PBS (**Fig. 1F**). This was presumably due to the phosphate affinity to metal sites, which caused the rapid breakdown of frameworks.^[31]^

The Fourier Transform Infrared (FT-IR) spectrum of Y-BTC (**Fig. 1G**) showed no peak near 1,700 cm^-1^, indicating the absence of free BTC molecules in the sample. Characteristic bands appear at 1,610 cm^-1^ and 1,540 cm^-1^, attributed to the higher-energy stretching vibrations of the C=O groups in the carboxylate moieties, while bands at 1,430 cm^-1^ and 1,360 cm^-1^ corresponded to the lower-energy bending vibrations of the same groups.^[25]^

The thermogravimetric analysis (TGA) of the Y-BTC MOFs was carried out to determine thermal stability and chemical composition. Following activation of Y-BTC at 150°C for 6 h to remove free and bound water molecules, its TGA curve (**Fig. 1H**) exhibited good thermal stability up to 400°C. Upon heating to approximately 600°C, Y-BTC was oxidated into Y_2_O_3_. The corresponding weight loss in the TGA curve was approximately 63.5 wt%, which is in close agreement with the theoretical value of 62.2 wt% for the activated Y-BTC.

Nitrogen porosimetry at 77K (**Fig. S3**) indicated that Y-BTC had a very small Brunauer-Emmett-Teller (BET) surface area of 10 m^2^ g^-1^, consistent with the value reported in previous literature.^[32]^

To assess retention of radioyttrium in MOFs, ^88^Y,Y-BTC MOFs were synthesized. Radioyttrium release profiles were measured in Milli-Q^®^ water, HEPES, PBS, and FBS (**Fig. 1I**). According to the obtained results, the produced particles demonstrated a strong yttrium retention with maximal loss of 5% ^88^Y found in FBS in 6 days at 37⁰C. Most likely, this is primarily caused by the fact that yttrium persists in precipitation as a component of YPO_4_ when phosphates are present that break the MOF particles. Thus, the long-term radioyttrium retention demonstrated by ^88^Y,Y-BTC rods, could be beneficial in the context of brachytherapy application.

### Synthesis and characterization of bimetallic MIL-100(Fe,Y) MOF particles

Bimetallic particles of MIL-100(Fe,Y) MOF (**Fig. 2A**) were synthesized following a previously reported eco-friendly method^[23]^ by mixing FeSO_4_·7H_2_O, Y(NO_3_)_3_·6H_2_O, trimesic acid, and NaOH in Milli-Q^®^ water at 60°C for 24 h, followed by centrifugation and washing with water and ethanol. Compared to the reference method,^[23]^ which implies room temperature, raising the temperature appropriately helps MIL-100 crystallize more uniformly, thus improving the homogeneous distribution of Y in the final product as well as significantly shorten the synthesis time (1 day at 60°C against 2 for the RT protocol). However, Y(BTC)(H_2_O)_6_ still appeared as a side product. As shown in **Fig. S4**, peaks at 13.5° and 17.4° in samples with n(Y)/[n(Y)+n(Fe)] ratios of 8% and 15% indicated the presence of traces of Y(BTC)(H_2_O)_6_.

**Figure 2.**
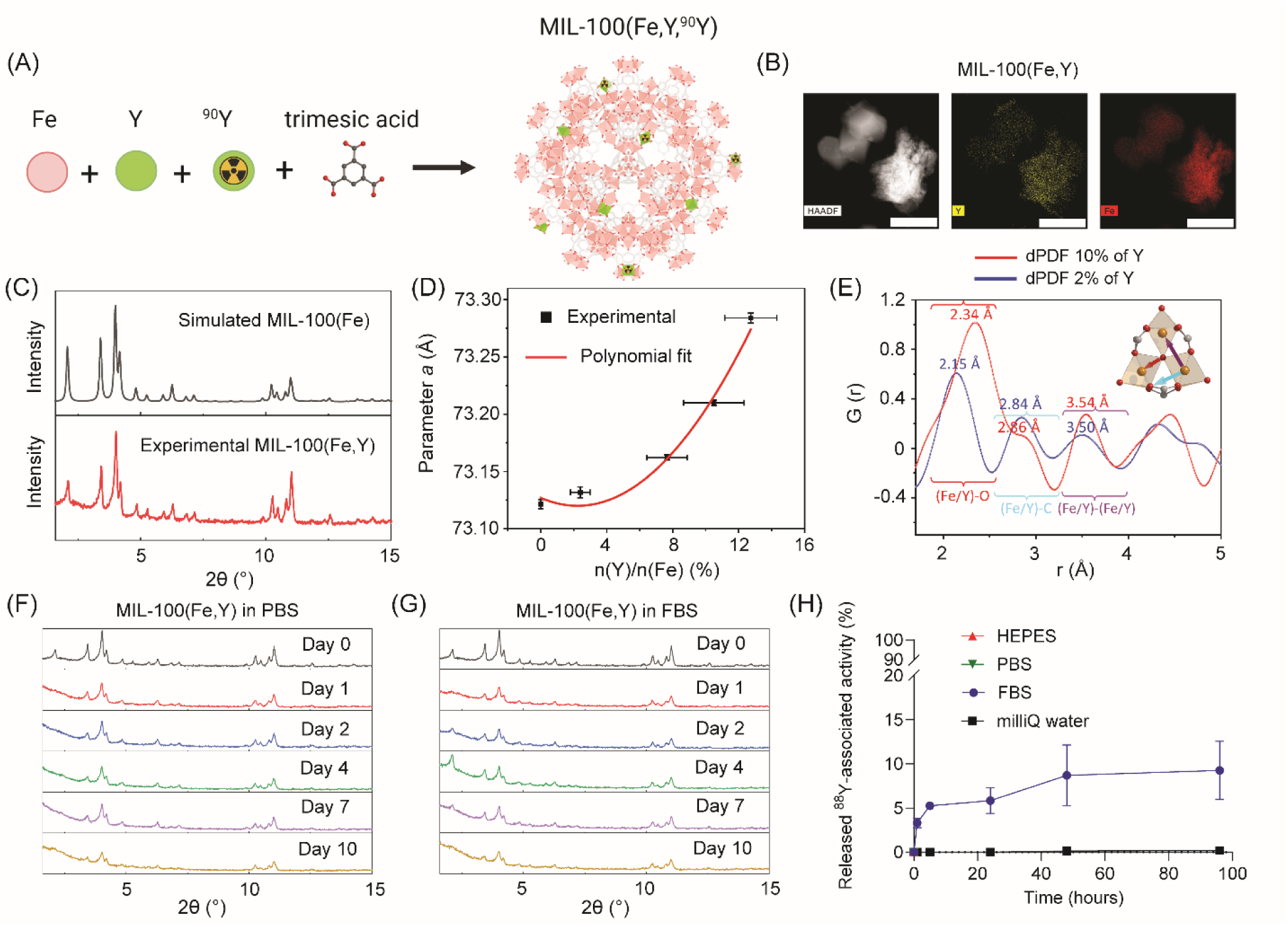
Physicochemical properties of MIL-100(Fe,Y) particles. (A) Scheme of MIL-100(Fe,Y) MOF structure after doping with radioyttrium. Oxygen atoms are shown in red, iron octahedra are shown in light red, yttrium octahedra are shown in light green. (B) STEM-HAADF EDX map of MIL-100(Fe,Y) particles with 10% of doped Y. The scale bar is 1 μm. (C) Powder X-ray diffraction patterns of simulated MIL-100(Fe) and experimental MIL-100(Fe,Y) (Co-K_α1_ radiation). (D) Non-linear dependence of the MIL-100(Fe) unit cell parameter (Mo-K_α1_ radiation) *a* on yttrium content (calculated by SEM-EDX). (E) Differential Pair Distribution Function (dPDF) analysis shows the elongation of the Fe/Y-O correlations upon the increase of Y content due to the appearance of Y-O bonds in the structure of MIL-100(Fe,Y). (F,G) PXRD patterns of MIL-100(Fe,Y) soaked in PBS (F) and FBS (G) over 0-10 days at 37°C. (H) *In vitro* yttrium-88 release from MIL-100(Fe,Y,^88^Y) MOFs in different solutions, including HEPES buffer, PBS, FBS, and purified MilliQ® water, at different time points (1, 3, 5, 24, 48, and 96 h) of incubation at 37⁰C.

Scanning Transmission Electron Microscopy – High Angle Annular Dark Field (STEM-HAADF) maps (**Fig. 2B**) and scanning electron microscopy coupled with energy-dispersive X-ray spectroscopy (SEM-EDX) analysis (**Fig. S5**, **Fig. S6**) confirmed the successful incorporation of Y into MIL-100 samples. In **Fig. S5**, a few rod-shaped microparticles were only found in samples with 8% Y, confirming the presence of a side product, Y(BTC)(H_2_O)_6_. For other samples with 2%, 5%, and 10% Y, they exhibited good homogeneity and purity (**Fig. 2B**, **Fig. S4**, and **Fig. S5**). SEM-EDX was also used to quantify the ratio of Y in resulting samples. As shown in **Table S1**, the values measured by SEM-EDX were similar to and even exceeded the amounts added during synthesis, indicating nearly 100% of Y loading efficiency. Thus, the 10% Y containing sample was selected as representative for further material characterization.

STEM-HAADF images (**Fig. 2B** and **Fig. S7**) showed that both MIL-100(Fe) and MIL-100(Fe,Y) exhibited a similar octahedral shape with aggregates of a few hundred nm. With their well-established rapid internalization *via* multiple endocytic pathways, MIL-100(Fe) nanoparticles are known to be highly effective vehicles for delivery of drugs to intracellular compartments.^[33,34]^ Dynamic Light Scattering (DLS) (**Fig. S8** and **Table S2**) indicated the hydrodynamic size of MIL-100(Fe) and MIL-100(Fe,Y) in PBS (pH=7.4) were both *ca.* 350 nm. However, their Polydispersity Index (PdI) was *ca.* 0.7, indicating a relatively broad particle size distribution. The zeta potential (**Table S2**) of MIL-100(Fe) and MIL-100(Fe,Y) were of -23.0±1.8 mV and - 24.3±1.3 mV, respectively. These values indicated that MIL-100(Fe) and MIL-100(Fe,Y) particles possessed similar negatively charged surfaces in PBS, suggesting a certain degree of electrostatic repulsion between particles and a moderate colloidal stability. The slight increase in surface charge with 10% of Y incorporation is here not significant, implying that the impact of Y doping on the surface charge of the particles is limited.

The crystal structure of the MIL-100(Fe,Y) was confirmed using PXRD as well as the differential pair distribution function (dPDF) analyses. The obtained powder diffraction patterns matched well with the known cubic crystal structure of MIL-100(Fe), confirming the formation of the MIL-100(Fe,Y) (**Fig. 2C**). In particular, the substitution of Fe atoms by Y led to a non-linear (polynom of the second order) increase of the cubic unit cell parameter *a* (**Fig. 2D**) due to the larger ionic radius of Y^3+^ compared to Fe^3+^. The complementary dPDFs showed that the doping of MIL-100(Fe) by Y was associated with much longer M-O bonds (2.34 Å for Y-O vs 2.0 Å for Fe-O), which were comparable with the ones known for its format (**Fig. 2E**).^[35]^

FT-IR analysis (**Fig. S9**) confirmed that both MIL-100(Fe) and MIL-100(Fe,Y) exhibited the main characteristic peaks of the reported MIL-100(Fe).^[33]^ The small peak at around 1,710 cm^-1^ was assigned to the stretching vibration of C=O groups, indicating residual traces of free trimesic acid in both samples.

TGA results (**Fig. S10**) showed that under heating in an oxygen atmosphere, both MIL-100(Fe) and MIL-100(Fe,Y) first experienced the removal of free and bound water molecules before 150°C, followed by the degradation of the constitutive ligand, corresponding to the weight loss of BTC. The residue at 600°C was attributed to iron oxides, or iron and yttrium oxides. As shown in **Fig. S10**, since yttrium is heavier than iron, the weight loss of BTC for MIL-100(Fe,Y) was slightly less than that of bare MIL-100(Fe).

Nitrogen porosimetry at 77K (**Fig. S11**) indicated that the BET surface area of MIL-100(Fe) and MIL-100(Fe,Y) were similar, of 1,570 m^2^ g^-1^ for MIL-100(Fe) and 1,650 m^2^ g^-1^ for MIL-100(Fe,Y). This suggested that doping with Y did not significantly change the availability of pores in MIL-100(Fe), thus the remaining surface area and pore volume were possible to encapsulate the potential therapeutic agents.

The stability of MIL-100(Fe,Y) under physiological conditions under physiological conditions was tested in PBS and FBS buffer. As shown in **Fig. 2F** and **Fig. 2G**, MIL-100(Fe, Y) exhibited good crystalline stability in both PBS and FBS buffer over a period of 10 days. These findings are consistent with some of our previous studies, indicating that MIL-100(Fe,Y) has good stability when the particle size is not too small.^[31],[36]^

To assess retention of radioyttrium in MIL-100(Fe,Y), MIL-100(Fe,Y,^88^Y) MOFs were synthesized. The obtained particles displayed almost no release of radioyttrium in Milli-Q® water, HEPES, and PBS (**Fig. 2H**). As in a case of Y-BTC MOFs, this phenomenon is rather attributed to yttrium precipitation as a part of YPO_4_ in the presence of phosphates. In FBS, a slow-rate radioyttrium release from MIL-100(Fe,Y,^88^Y) was found, with roughly 10% release in 120 h at 37⁰C. Noteworthy, the release studies demonstrated that these materials retain radioyttrium at a high rate for a long period indicating their promise for brachytherapy.

### Tumor retention of radioyttrium upon intratumoral administration of ^88^Y-doped MOFs

The ability of the carrier to retain radionuclides at the tumor site is crucial for the effectiveness of intratumoral radiotherapy. We conducted an *in vivo* study to assess the retention of radioactivity in the tumor after the local injection of radioyttrium-doped MOFs. Since yttrium-90 is a pure beta-emitter with a short half-life, we utilized our MOFs doped with yttrium-88, a gamma-emitter (898 keV and 1,836 keV) with a half-life of 106.6 days,^[37]^ making it more suitable for radionuclide biodistribution measurement.

MIL-100(Fe,Y,^88^Y) particles were administered intratumorally to LLC1 tumor-bearing mice, and the ^88^Y radioactivity in the tumor and main organs (liver, kidneys, lungs, heart, bladder, and intestines) was measured at several time points (1, 3, and 7 days) (**Fig. 3A**) and determined as a percentage of the injected dose per gram of tissue (% ID/g) (**Fig. 3B**) or per organ (% ID) (**Fig. 3C**). On day 1, approximately 80% ID accumulated in the tumor, while a small percentage of ID was detected in the liver and in the body (likely in blood) that could be a result of a minor leakage of MIL-100(Fe,Y,^88^Y) particles to the circulation. By day 3, we still observed 80% ID remaining in the tumor, indicating no significant washout of ^88^Y from the tumor site during this period (**Fig. 3C**). However, by one week post-injection, the majority of radioactivity had been cleared from the tumor and had accumulated in the body and in the liver. This suggests a rapid biodegradation of MIL-100(Fe,Y,^88^Y) particles after day 3. Considering that at least 80% ID remains in the tumor tissue for 3 days, a simple approximation for ^90^Y (see Supplementary Methods) suggests that at least 38% of decays will occur in the tumor. Despite the relatively high washout of radioyttrium from the tumor after day 3, this does not result in a significant increase in the percentage of ID per gram of tissue in other organs (**Fig. 3B**), thus minimizing the radioactive exposure of off-target tissues.

**Figure 3.**
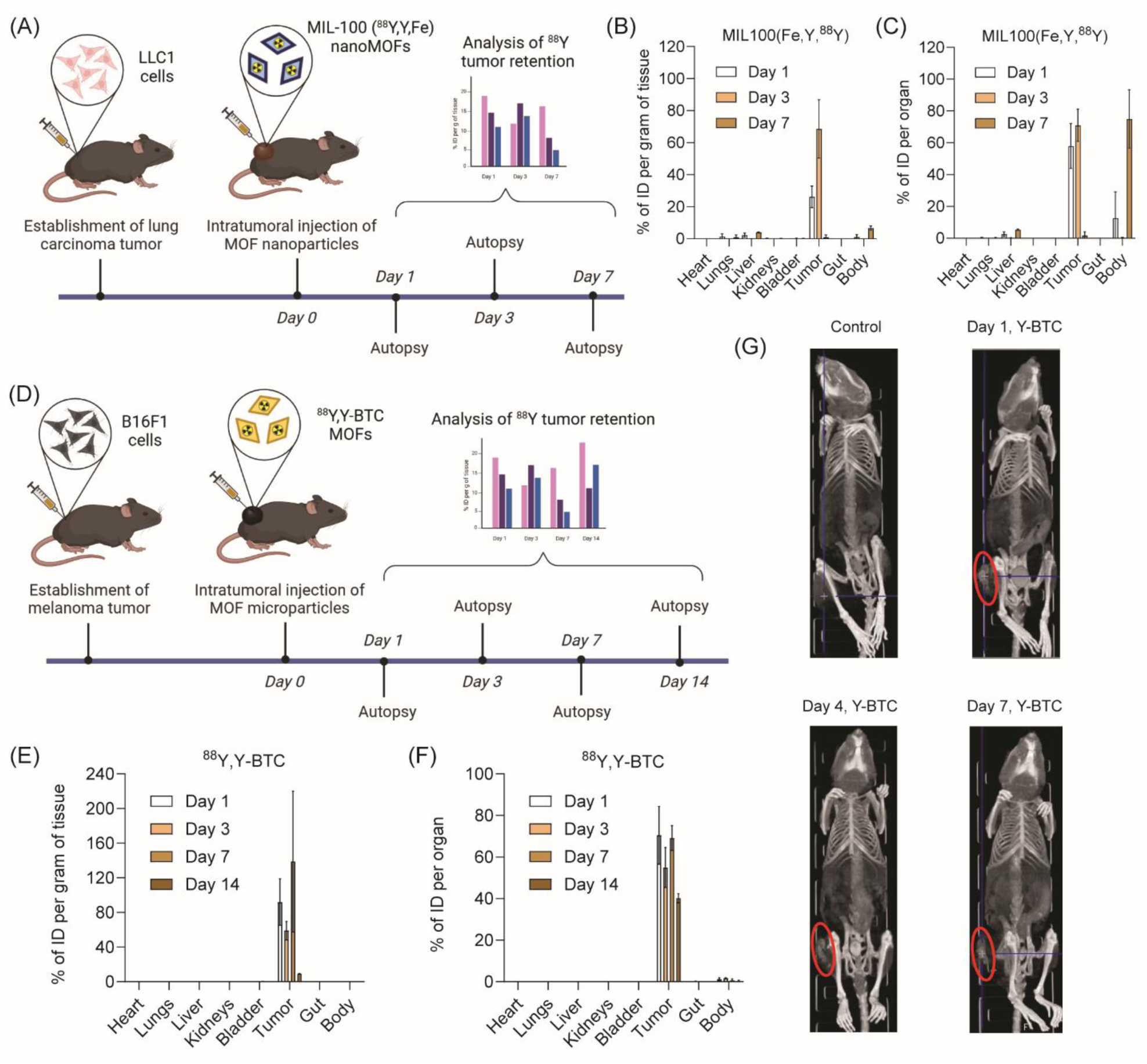
Yttrium-88 and MOF retention at tumor site after intratumoral injection of ^88^Y-doped MOFs. (A) A scheme of the experiment with LLC1 tumor-bearing mice treated with MIL-100(Fe,Y,^88^Y) particles at a single dose of 1.25 kBq (0.034 μCi). (B) Radioactivity measurements in different tissues on days 1, 3, and 7 after treatment with MIL-100(Fe,Y,^88^Y) particles shown as the percentage of injected dose (ID) per gram of tissue and (C) per organ. The data are expressed as the mean ± SD (n = 3). (D) A scheme of the experiment with B16F1 tumor-bearing mice treated with ^88^Y,Y-BTC particles at a single dose of 1.4 kBq (0.038 μCi). (E) Radioactivity measurements in different tissues on days 1, 3, 7, and 14 after treatment with ^88^Y,Y-BTC particles shown as the percentage of ID per gram of tissue and (F) per organ. The data are expressed as the mean ± SD (n = 3). (G) Reconstructed 3D-projections of mice on days 1, 4, and 7 after intratumoral injection of Y-BTC particles versus a control mouse (injected with saline). Red ellipse outlines the contrast area at the site of MOF administration.

Since Y,^88^Y-BTC MOF exhibited stronger retention of radioyttrium in FBS than MIL-100(Fe,Y,^88^Y), we conducted measurements of ^88^Y radioactivity in the tumor and other organs over a two-week period with four different time points at 1, 3, 7, and 14 days after intratumoral injection of Y,^88^Y-BTC particles in B16F1 cell tumor-bearing mice (**Fig. 3D**). As previously shown, tissue radioactivity was determined as a percentage of the injected dose per gram of tissue (% ID/g) (**Fig. 3E**) or per organ (% ID) (**Fig. 3F**). It was observed that Y,^88^Y-BTC particles demonstrate robust retention of ^88^Y radioactivity in the tumor, reaching 80% ID by the first week and 40% by 14 days after injection. This retention profile is particularly advantageous for the therapeutic efficacy of brachytherapy. Considering this result, a simple approximation for ^90^Y (see Supplementary Methods) suggests that at least 75% of decays will occur in the tumor. To emphasize the benefits of yttrium incorporation into MOFs and for the sake of comparison, we evaluated the retention of intratumorally injected yttrium-88 in the form of yttrium chloride (YCl_3_) on day 1. It was observed that the intratumoral activity of ^88^YCl_3_ dropped two-fold more strongly compared to ^88^Y,Y-BTC, while also exhibiting enhanced off-target accumulation in the kidneys, liver, and body (**Fig. S12**). Importantly, there was no noticeable accumulation of ^88^Y radioactivity beyond the tumor after intratumoral injection of Y,^88^Y-BTC particles (**Fig. 3E** and **Fig. 3F**). Since the yttrium-containing MOFs demonstrated sufficient contrast efficacy in X-ray computed tomography (CT), we obtained their images in the tumor. Consistent with tissue radioactivity measurements, the injected particle depot remained at the site of injection for at least one week (**Fig. 3G**). Hence, the capability of our MOFs to be visualized by CT is crucial for ensuring uniform tumor exposure to radioactivity, which presents an advantage over polymer-based biodegradable radionuclide carriers.

The obtained results have shown that the Y-BTC particles exhibited better radioyttrium retention in the tumor compared to MIL-100(Fe,Y), which could be attributed to their slower biodegradation behavior. Such a difference in stability might be due to the differences in particle size and porosity. Overall, Y-BTC particles demonstrated sufficient retention time for ^90^Y to achieve maximal radiation damage to the tumor while reducing harm to normal tissue.

### Uptake and growth inhibition of the A549 tumor spheroids by ^90^Y-doped MIL-100(Fe,Y) particles

As opposed to conventional 2D cell cultures, multicellular spheroids display characteristics reminiscent of tumor microenvironments, such as the development of nutrient gradients, a hypoxic core, abundant cell-cell interactions, and the secretion of extracellular matrix (ECM) by spheroid cells. Moreover, 3D tumor spheroids better mimic the release of soluble bioactive molecules, gene expression profiles, and drug resistance.^[38]^ For this reason, they are often employed as a predictive platform to evaluate new drug candidates. To our knowledge, tumor spheroids have not yet been used for testing of radioyttrium formulations.

Once MIL-100(Fe,Y) particles exhibit a mean diameter in the range of 300-400 nm, they could be internalized by cancer cells as opposed to large Y-BTC particles. To estimate this, MIL-100(Fe,Y) particles were fluorescently labeled with TBO dye and then added to 3D A549 spheroids. Particle uptake and penetration were evaluated in DAPI-stained spheroid sections using fluorescent microscopy after 12 and 24 h post-treatment (**Fig. 4A**). At the initial time point of 12 h, the fluorescent MIL-100(Fe,Y) particles accumulated in the outer cell layers of the spheroids. Subsequently, after 24 h of incubation, a greater number of particles were observed to have penetrated the tumor spheroids (**Fig. 4A**). This indicates that the particles are capable of entering the 3D A549 spheroids and gaining access to the central region of the tumor to some extent. It is noteworthy that a significant portion of the MIL-100(Fe,Y) particles were co-localized with DAPI (the Mander’s overlap coefficients for the TBO (M1) and the DAPI (M2) channels are 0.7103 and 0.9974, respectively), indicating mainly intracellular deposition of the particles within the spheroids (**Fig. 4B**). Analysis of fluorescence intensity profiles has shown that fluorescently tagged particles readily crossed up to a 50-micron distance from the margin (**Fig. 4C**), corresponding to three to five cell layers. Particle uptake likely contributes to reaching the spheroid core to some extent as was shown earlier.^[39]^ Thus, MIL-100(Fe,Y) particles undergo cellular uptake and penetrate tumor spheroids.

**Figure 4.**
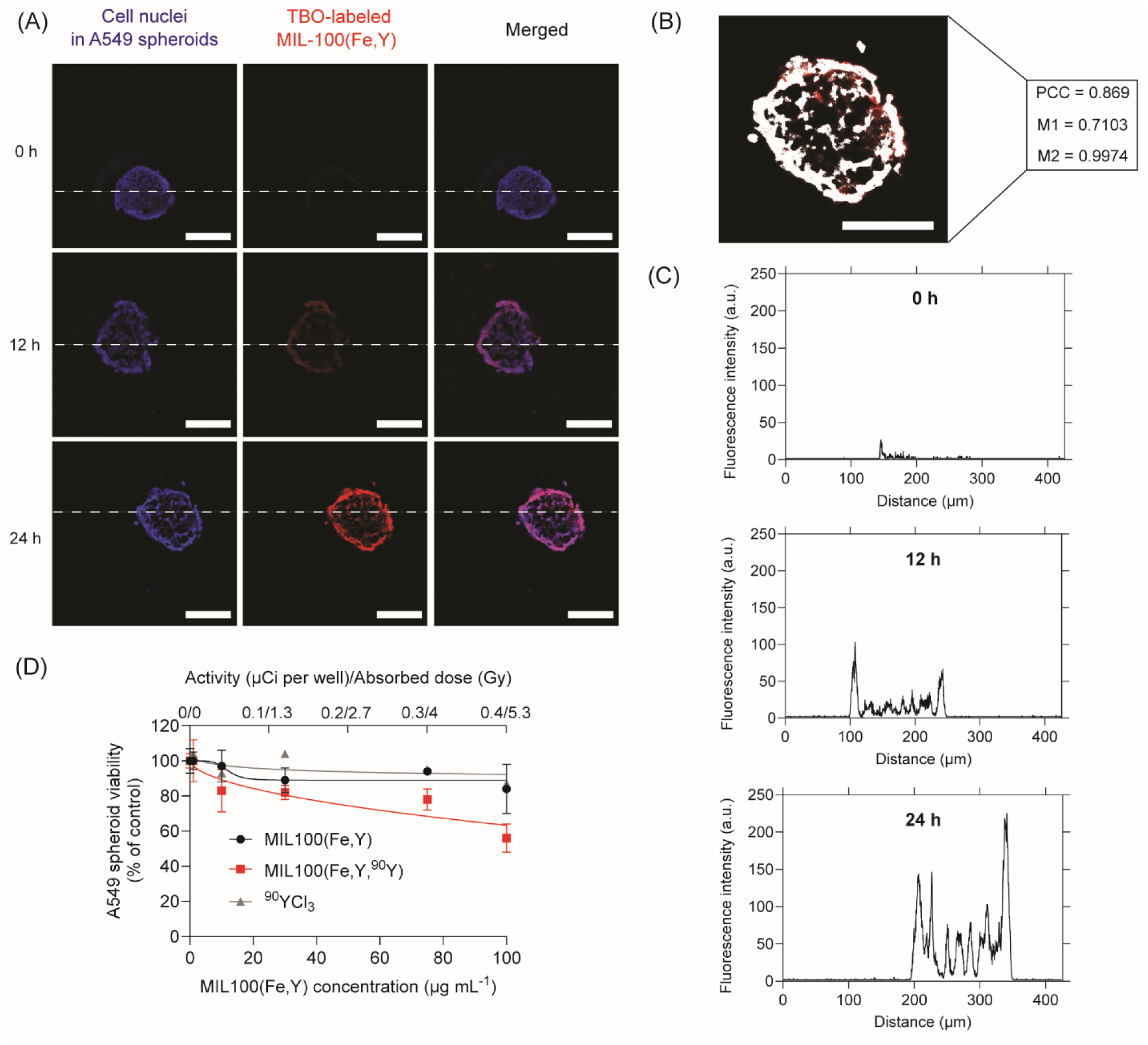
Uptake and growth inhibition of A549 tumor spheroids induced by ^90^Y-doped MIL-100(Fe,Y) particles. (A) The representative confocal images of A549 spheroids after incubation with fluorescently TBO-labeled MIL-100(Fe,Y) particles (red) at the concentration of 100 µg mL^-1^ for 12 and 24 h. Staining with DAPI (blue) determines the cell nuclei. The scale bar is 100 µm. (B) The representative image of the spheroid section after 24-h incubation with TBO-labeled MIL-100(Fe,Y) particles in TBO channel, where the white fraction of pixels co-localizes with the DAPI channel. The values of the Pearson correlation coefficient (PCC) and the Mander’s overlap coefficients for the TBO (M1) and the DAPI (M2) channels are provided. The scale bar is 100 µm. (C) Fluorescence intensity profiles in the TBO channel along the dashed horizontal line from (A) displaying penetration of TBO-labeled MIL-100(Fe,Y) MOF particles to A549 spheroids. (D) Spheroid viability on day 5 after treatment with ^90^YCl_3_, MIL-100(Fe,Y), and MIL-100(Fe,Y,^90^Y) at different concentrations. The values are shown as % of control (non-treated A549 spheroids). The data are expressed as the mean ± SD (n = 5).

To assess the antitumor effects of MIL-100(Fe,Y,^90^Y), MIL-100(Fe,Y), or ^90^YCl_3_, the spheroids were incubated with the formulations for five days, followed by cell viability assay based on ATP level measurement (see Materials and methods). It has been shown that neither ^90^YCl_3_, nor MIL-100(Fe,Y) particles have not caused cytotoxic effects at concentrations ranging from 0.004 to 0.4 µCi per well and from 1 to 100 µg mL^-1^, respectively. In contrast, cytotoxicity of MIL-100(Fe,Y,^90^Y) particles increased in a dose-dependent manner with fewer than 60% of survived cells after being exposed to MIL-100(Fe,Y,^90^Y) at a concentration of 100 µg mL^-1^ (or 0.4 µCi per well) (**Fig. 4D**). It should be noted that the effect of cellular uptake of beta-emitters on the cell viability is limited because of the relatively long range of beta particles. However, the observed effect suggests that the MIL-100(Fe,Y,^90^Y) particles are capable of enhancing beta-emitted cytotoxicity.

Based on the obtained data, it is challenging to assess the benefit of cellular uptake of MIL-100(Fe,Y,^90^Y) particles. On one hand, it sensitizes the cytotoxic effects of radioyttrium. On the other hand, it could lead to accelerated decomposition of the MOFs in the endosome/lysosome compartment and subsequent washout of radionuclides from the tumor.

### Growth inhibition of the B16F1 spheroids by ^90^Y

The cytotoxicity of yttrium-90 was assessed in B16F1 cell spheroids to determine the minimum therapeutic activity dose for *in vivo* experiments. Growth curves of B16F1 spheroids treated with various doses of ^90^YCl_3_ were obtained by microscopic measurement of spheroid volume (**Fig. 5**).

**Figure 5.**
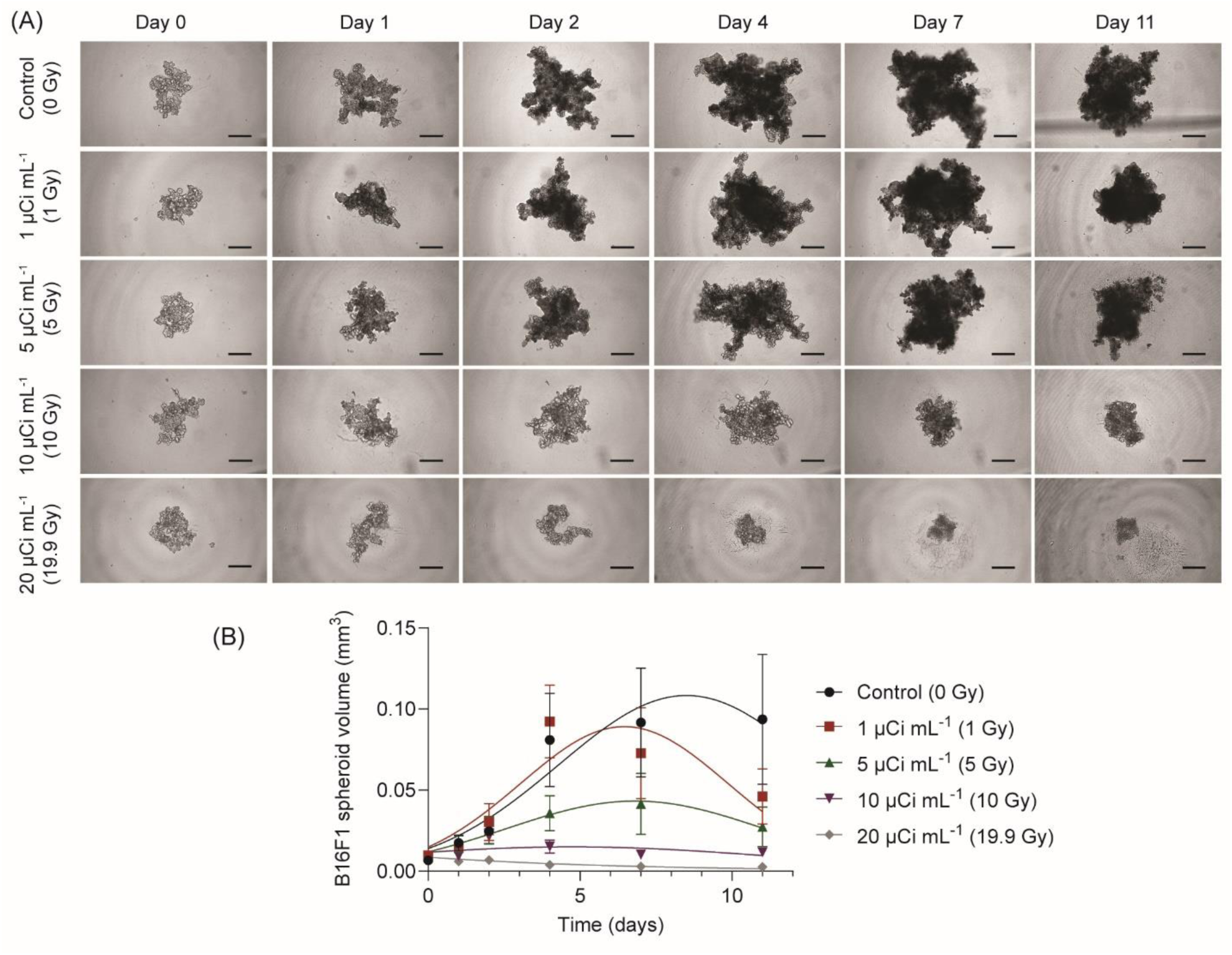
Determination of the minimal therapeutic dose of yttrium-90 using the B16F1 spheroid model. (A) Representative images of the spheroids after treatment with different doses of ^90^YCl_3_ (0, 1, 5, 10, and 20 μCi mL^-1^). The scale bar is 200 μm. (B) Growth curves of B16F1 spheroids following the treatment with different doses of ^90^YCl_3_. The data are expressed as the mean ± SD (n = 5).

The control spheroids (non-treated) exhibited typical growth behavior, which can be described using a Gompertz function (**Fig. 5A**). The asymptotic volume of approximately 0.1 mm^3^ was achieved by the control spheroids by day 4 and remained unchanged by day 11. Spheroids treated with yttrium-90 displayed similar growth kinetics. As expected, we observed a dose-dependent inhibition of spheroid growth rate (**Fig. 5B**). At a dose of 20 μCi mL^-1^, corresponding to 19.9 Gy, the spheroids have not demonstrated any growth. Therefore, we considered this dose as the minimum therapeutic dose for further *in vivo* experiments.

### Antitumor efficacy of the ^90^Y,Y-BTC MOFs

The radiotherapy efficacy of the ^90^Y,Y-BTC MOFs was tested using C57BL/6J mice bearing B16F1 tumors. The mice with the average tumor volume of 149±38 μL were randomly divided into three groups (8 mice per group). The mice in the treatment group (group 1) received ^90^Y,Y-BTC particles at a radioactivity dose of 0.25 μCi per mm^3^ of tumor, with an injection concentration of 100 mg per mL Y-BTC. The injection volume with MOF particles, equivalent to one-fourth of the tumor size, was delivered as a single, bolus infusion into the tumor core. Control mice were injected with unlabeled Y-BTC microparticles at the same concentration (group 2) or saline (group 3) at the same volume. It should be noted that specific activity of ^90^Y,Y-BTC MOFs is tunable. In this study, it was approximately 5×10^-5^ % of maximal, which provides an opportunity to vary it depending on required injection volume and MOF concentration. The mice were subjected to euthanasia when reaching a tumor volume above 1 cm^3^.

It has been shown that brachytherapy with ^90^Y-doped Y-BTC MOFs impeded tumor growth (**Fig. 6A**) and extended the survival time of mice compared to the control group administered with saline (**Fig. 6B**). In the saline control group, 50% of the mice had tumor volumes surpassing 1 cm^3^ by day 4 post-treatment (median survival of 4.5 days). By day 7, this percentage had reached 100 %. In the group receiving treatment with ^90^Y,Y-BTC, the administration of radioactive elements was seen to effectively inhibit tumor growth and prolong median survival time up to 8.5 days. Another control group treated with non-radioactive Y-BTC particles also exhibited impeded tumor growth and prolonged median survival (6.5 days), although these differences were not statistically significant as compared with the saline-treated group. Observed effects may be attributed to cytotoxic effects of the particles on tumor cells.

**Figure 6.**
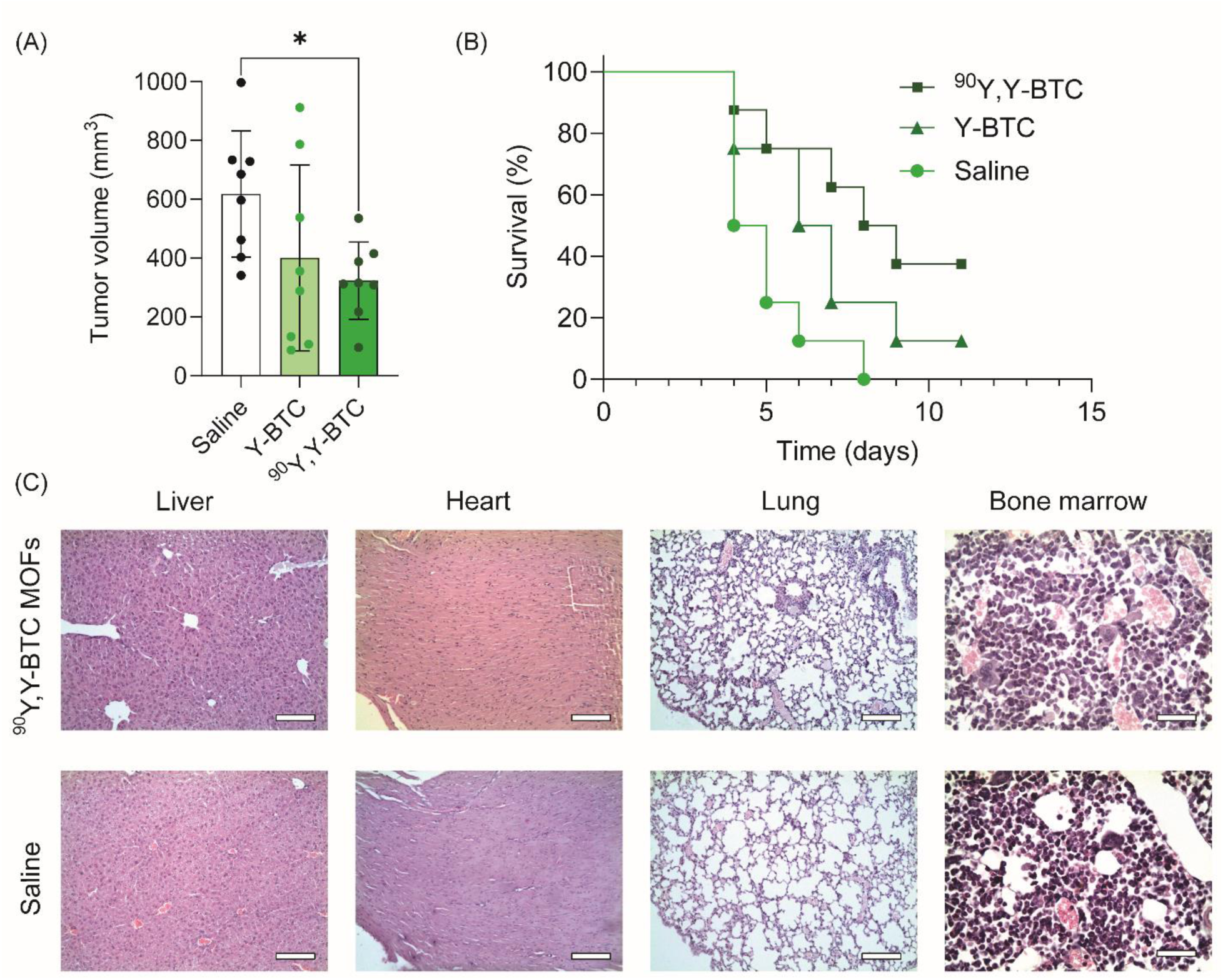
*In vivo* therapeutic efficacy of ^90^Y,Y-BTC MOFs after intratumoral administration. (A) Mean tumor volumes on day 3 after the treatment. The data are expressed as the mean ± SD (n = 8). *p < 0.05 (one-way ANOVA followed by a *post hoc* Dunnett’s test). (B) The survival curves of the experimental groups treated with saline, Y-BTC particles, and ^90^Y,Y-BTC particles (9.2 MBq per mL of tumor volume). *p < 0.05 (survival differences were determined by log-rank Mantel-Cox test). (C) Histological images of different tissue sections, demonstrating lack of histopathological changes. Scale bar is 100 μm.

Regarding potential side effects, mice injected with ^90^Y,Y-BTC exhibited no appreciable drop in body weight and displayed no abnormal behaviors after treatment. Histological examination of organ slices stained with H&E revealed no notable signs of organ damage in mice that survived at the endpoint after receiving a single dose of ^90^Y,Y-BTC particles (**Fig. 6C**).

## Discussion

Yttrium-90, a high-energy beta-emitter (up to 2.27 MeV) with a half-life of 64.1 h, is a widely used radioisotope in different brachytherapeutic modalities including hepatic radioembolization with ^90^Y glass microspheres for treatment of liver metastasis,^[40]^ brachytherapy of solid tumors in veterinary medicine by intratumoral injection of ^90^Y-doped yttrium phosphate particles in gel formulation,^[41,42]^ episcleral treatment of ocular malignancies using ^90^Y beta-emitting titanium sources,^[43]^ and radiosynovectomy by intra-articular injection of ^90^Y hydroxyapatite colloid.^[44]^ Yttrium-90 beta particles penetrate tissues to a maximum depth of approximately 11 mm, making it an ideal brachytherapy agent for larger tumors. High-energy beta-particles hold promise for effectively irradiating tumor tissue, compensating for minor source distribution inhomogeneities.^[45]^

Here, we fabricated two different yttrium-90-doped MOFs for nuclear medicine applications into a single step. Our method allowed us to incorporate yttrium into MOFs with different structure and porosity. Radioyttrium was incorporated into Y-BTC and MIL-100 MOFs. While doping of yttrium is predictable for yttrium-based Y-BTC, it was crucial to confirm it for the iron-based MIL-100 structure, which has been extensively used in multiple *in vivo* studies for the delivery of anticancer drugs.^[15,46,47]^ Noteworthy, the substitution of Fe atoms by Y was confirmed as shown by the increase in unit cell volume and the formation of new Y-O bonds **(Fig. 2D and Fig. 2E**). Importantly, iron was uniformly incorporated to MIL-100 structure according to STEM-HAADF EDX data (**Fig. 2B**).

The analysis of yttrium-88 release from MIL-100(Fe,Y,^88^Y) and ^88^Y,Y-BTC particles indicated a slow release profile of the radioisotope. In MIL-100(Fe,Y,^88^Y), release was higher in blood serum (**Fig. 2H**), presumably due to the faster degradation of MIL-100 in the presence of phosphates or citrate from the serum. It cannot be excluded that a part of the radioyttrium was transferred to non-soluble yttrium oxide or yttrium phosphate. In contrast, ^88^Y,Y-BTC released a very small amount of yttrium-88 for 6 days (**Fig. 1I**). The observed result could be attributed to the larger particle size compared to MIL-100 particles. Additionally, the low release of ^88^Y could be explained by surface degradation of Y-BTC MOF structure and its conversion into non-soluble yttrium oxide or yttrium phosphate. Alternatively, the formation of yttrium oxide or yttrium phosphate shell on the particle surface could delay degradation of Y-BTC MOF structure and further ^88^Y release.

The analysis of radioactivity retention in the tumor after intratumoral administration of ^88^Y,Y-BTC confirmed the formation of a stable radioactive depot. Based on the data for ^88^Y-doped Y-BTC MOFs (with an ^88^Y half-life of 106.6 days), a rough approximation for ^90^Y,Y-BTC shows at least 75% of localized activity deposition after two weeks. The observed radioisotope retention rate for Y-BTC particles was comparable to or superior to that of other biodegradable polymeric and MOF carriers. For example, ^131^I-modified polypeptide cross-linked hydrogels exhibited 60-80% radioisotope tumor retention after two weeks post-injection,^[8]^ while Y-BTC MOFs demonstrated 40% of radioyttrium retention after the same term (**Fig. 3F**). When considering MOF-based systems, only two studies provide available radioisotope biodistribution data after intratumoral injection. In the first study, data were obtained for polyethylene glycol (PEG)-modified nanoMOFs composed of hafnium (Hf^4+^) and tetrakis (4-carboxyphenyl) porphyrin (TCPP), where the porphyrin structure was exploited for coordination of ^99m^Tc^4+^. It was found that the % ID dropped from 100% to 20% for 24 h.^[18]^ Another study, which utilized ^177^Lu-doped MIL-101(Fe) decorated with PEG-folate acid conjugates for brachytherapy, provided only short-term imaging data (48 h post-injection), showing no significant decrease in activity within the tumor. It should be noted that MIL-101(Fe) MOFs has a close composition and structure to MIL-100 studied here leading to a faster degradation in body fluids. In turn, MIL-100(Fe,Y,^88^Y) exhibited extensive radioisotope washout from the tumor after day 3 (**Fig. 3B** and **Fig. 3C**).

Considering radioyttrium washout from the tumor, the tissues displaying the highest off-target activity were the liver and the body carcass (**Fig. 3B** and **Fig. 3C**). Our data are consistent with some early studies,^[48,49]^ which have shown that intraperitoneally or intravenously administered yttrium in yttrium chloride form is excreted from the circulation *via* the hepatobiliary rather than renal pathway. In addition, some accumulation in bones has been observed.^[48,49]^ It is noteworthy that the values of off-target yttrium accumulation for Y-BTC particles were insignificant (less than 1.5% ID). Importantly, the fact of yttrium washout from Y-BTC depots indirectly indicates a slow-rate *in vivo* decomposition of these materials. As for MIL-100(Fe,Y,^88^Y) particles, they demonstrated at least a 3-day tumor retention, followed by robust washout (**Fig. 3C**). We expect that upon surface modification of the external surface of MIL-100 nanoparticles, the resulting protective shell could slow down their decomposition and increase radioyttrium retention at the depot.^[50]^

Synthesis of particles with increased size could also lead to a slower degradation of MIL-100(Fe,Y) and radioyttrium release. Alternatively, considering the ability of MIL-100 particles for cellular uptake, one could consider this MOF for intracellular delivery of shorter living alpha-emitters such as ^213^Bi or ^211^At.

Anticancer efficacy of MIL-100(Fe,Y,^90^Y) was evaluated using A549 lung tumor spheroids. It has been shown that MIL-100(Fe,Y) particles were able to penetrate the spheroids within 24 h likely due to transcytosis rather than paracellular diffusion because of high co-localization extent with cell nuclei (**Fig. 4A** and **Fig. 4B**). Treatment of the spheroids with ^90^Y-doped MIL-100(Fe,Y) resulted in superior cytotoxic effects compared to non-radioactive MIL-100(Fe,Y) particles added at the same concentration. Interestingly, the same activity of ^90^YCl_3_ added to the spheroids did not cause significant cytotoxicity (**Fig. 4D**). The increased toxicity of ^90^Y-doped MIL-100(Fe,Y) MOFs compared to ^90^YCl_3_ may be attributed to the ability of MIL-100(Fe) to sensitize ferroptosis, a form of regulated cell death characterized by iron-dependent lipid peroxidation. It has been shown that the Fe^3+^ ions delivered by MIL-100(Fe,Y) could be reduced and then serve as potent inducers of ferroptosis by initiating the Fenton reaction, thereby promoting oxidative stress in cancer cells.^[51]^ As a result, the combined effect of yttrium-90 beta-decay and MIL-100-induced ferroptosis led to almost 50% decrease in cell viability in the A549 spheroids (**Fig. 4D**).

The anticancer effect of ^90^Y,Y-BTC particles was evaluated using a tumor-bearing mouse model. To establish the minimal therapeutic dose of yttrium-90, we employed a melanoma tumor spheroid model, as spheroids typically exhibit higher drug resistance rates than cell monolayers due to impaired drug penetration, increased pro-survival signaling, and hypoxic conditions.^[29,52,53]^ Earlier studies evaluating the *in vitro* anticancer effects of yttrium-90 and its immunoconjugates have utilized colony formation assays^[54]^ or metabolic viability assays.^[55]^ In contrast to these assays, daily measurement of tumor spheroid diameters allows for real-time determination of growth inhibition rates. It was determined that the IC50 dose for B16F1 spheroids on day 11 ranged between 5 and 10 Gy (**Fig. 5**). To compensate for the extensive growth rate of B16F1 tumors in further *in vivo* therapeutic experiments, we used an almost 20-fold higher dose. The selected dose of ^90^Y,Y-BTC particles (50 μCi per 200 mm^3^ tumor corresponding to around 380 Gy for a week exposure) significantly delayed tumor growth and prolonged survival rate compared to the control group (**Fig. 6A** and **Fig. 6B**). Although not statistically significant, treatment with Y-BTC particles also caused some delay in tumor growth (**Fig. 6A**) that could be attributed to previously reported cytotoxic effects of yttrium including overproduction of reactive oxygen species (ROS), decrease in ATP production and DNA damage.^[56]^

Noteworthy, treatment with ^90^Y,Y-BTC particles did not cause any histopathological changes in the tissues beyond the tumor including the most radioresistant bone marrow (**Fig. 6C**). Thus, the insignificant off-target accumulation of radioactivity, combined with the absence of systemic side effects confirmed by histological analysis, implies that any MOF components that escape the tumor are safely eliminated from the body.

## Conclusions

In summary, we have developed biodegradable metal-organic frameworks based on bimetallic MIL-100(Fe,Y) and Y-BTC particles doped with yttrium-90 for low-dose rate intratumoral radiotherapy. Both MOF particles have been shown to retain yttrium in different buffer solutions, as well as to retain radioyttrium at the tumor site after intratumoral injection, without significant accumulation of yttrium in off-target tissues. CT contrasting abilities of MOFs allow to visualize tumor coverage with radioactive particles that distinguishes them favorably from polymeric biodegradable carriers for brachytherapy. After intratumoral injection, ^90^Y-doped Y-BTC MOFs effectively inhibited the growth of B16F1 melanoma tumors and prolonged the survival of tumor-bearing mice. Therefore, our research demonstrates the possibility of creating biodegradable and non-toxic MOF particles carrying radioactive ^90^Y, which is able to provide a valuable strategy for further application in low-dose rate internal radiation treatment. Notably, MOFs demonstrate an almost limitless capability to vary the specific activity of the injectable material over a wide range. Although we did not take here the benefit of drug encapsulation into ^90^Y-doped MOF structures, the feasibility of generating MOFs with varying porosities is important for additional encapsulating therapeutic molecules of different sizes to achieve a combined chemoradiotherapeutic effect and/or be combined with photosensitizers, photothermal agents, proteins, and antibodies, enabling future research to investigate synergistic combinations of treatments.

## Acknowledgements

The authors acknowledge the Ministry of Science and Higher Education of the Russian Federation for financial support (agreement № 075-15-2021-1363). This work was also supported by the Ministry of Science and Higher Education of the Russian Federation within the framework of state support for the creation and development of a World-Class Research Center “Digital Biodesign and Personalized Healthcare”, project № 075–15-2022–306. The authors are thankful to Dr. Alexander Masyutin and Dr. Maria Erokhina from Lomonosov Moscow State University for their help with histological analysis of tissue samples. TG-DTA measurements were performed using the equipment of the Center of shared use of IPCE RAS. Experiments with CT-imaging were carried out with the use of equipment from Lomonosov Moscow State University Program of Development.

