## Supplementary Data for "Yttrium-90-doped metal-organic frameworks (MOFs) for low-dose rate intratumoral radiotherapy"

### Supplementary Figures

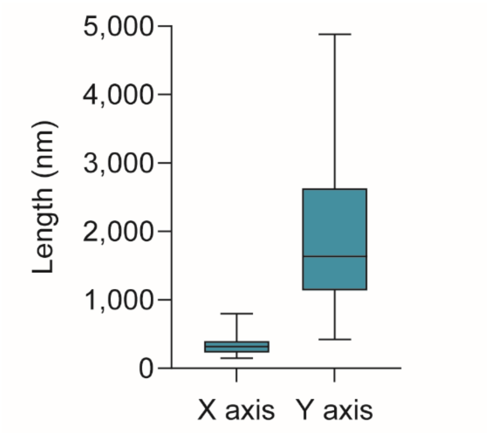


**Figure S1.** Size distribution of Y-BTC miroparticles for X and Y axis determined by quantitative analysis of SEM images. Over 30 particles were measured. Approximate aspect ratio is 1:6. The data are expressed as the mean ± SD.

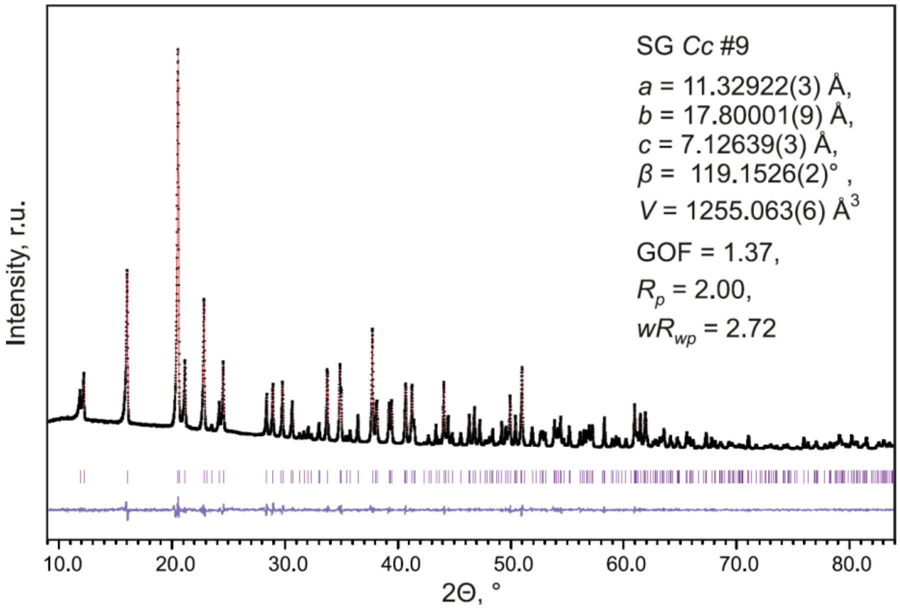


**Figure S2.** Powder X-ray diffraction pattern of experimental Y-BTC particles (Co-$K_{\alpha 1}$ radiation).

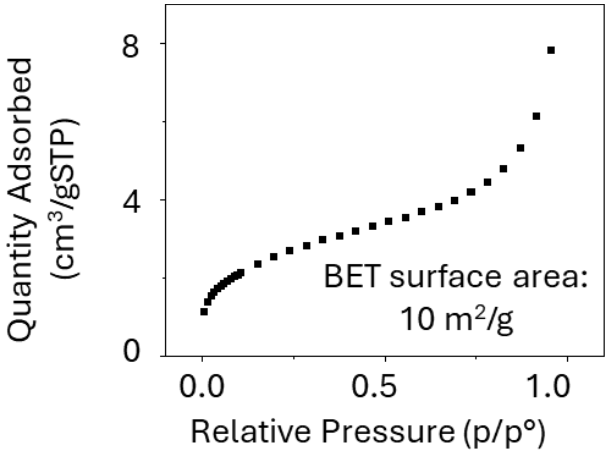


**Figure S3.** Nitrogen adsorption analysis of Y-BTC.

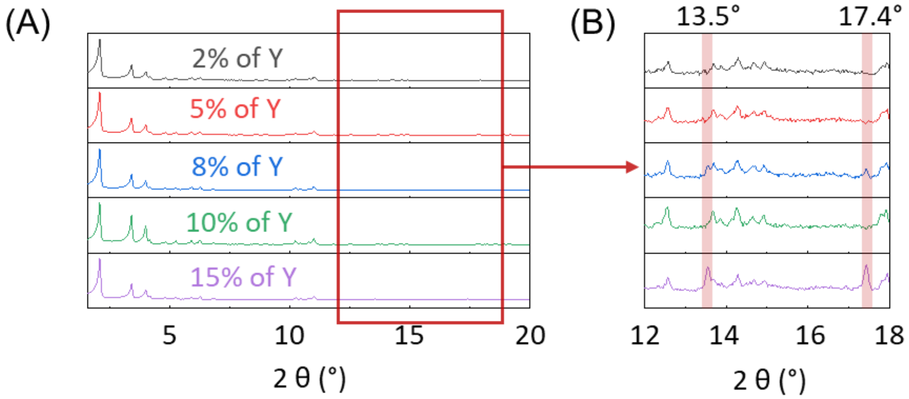


**Figure S4.** (A) PXRD pattern of MIL-100(Fe,Y) with n(Y)/[n(Y)+n(Fe)] ratios ranging from 2% to 15%. (B) Magnified PXRD pattern focusing on the range from 12 to 18°, highlighting the peaks at 13.5° and 17.4°, which correspond to the presence of Y(BTC)$(H_2O)_6$.

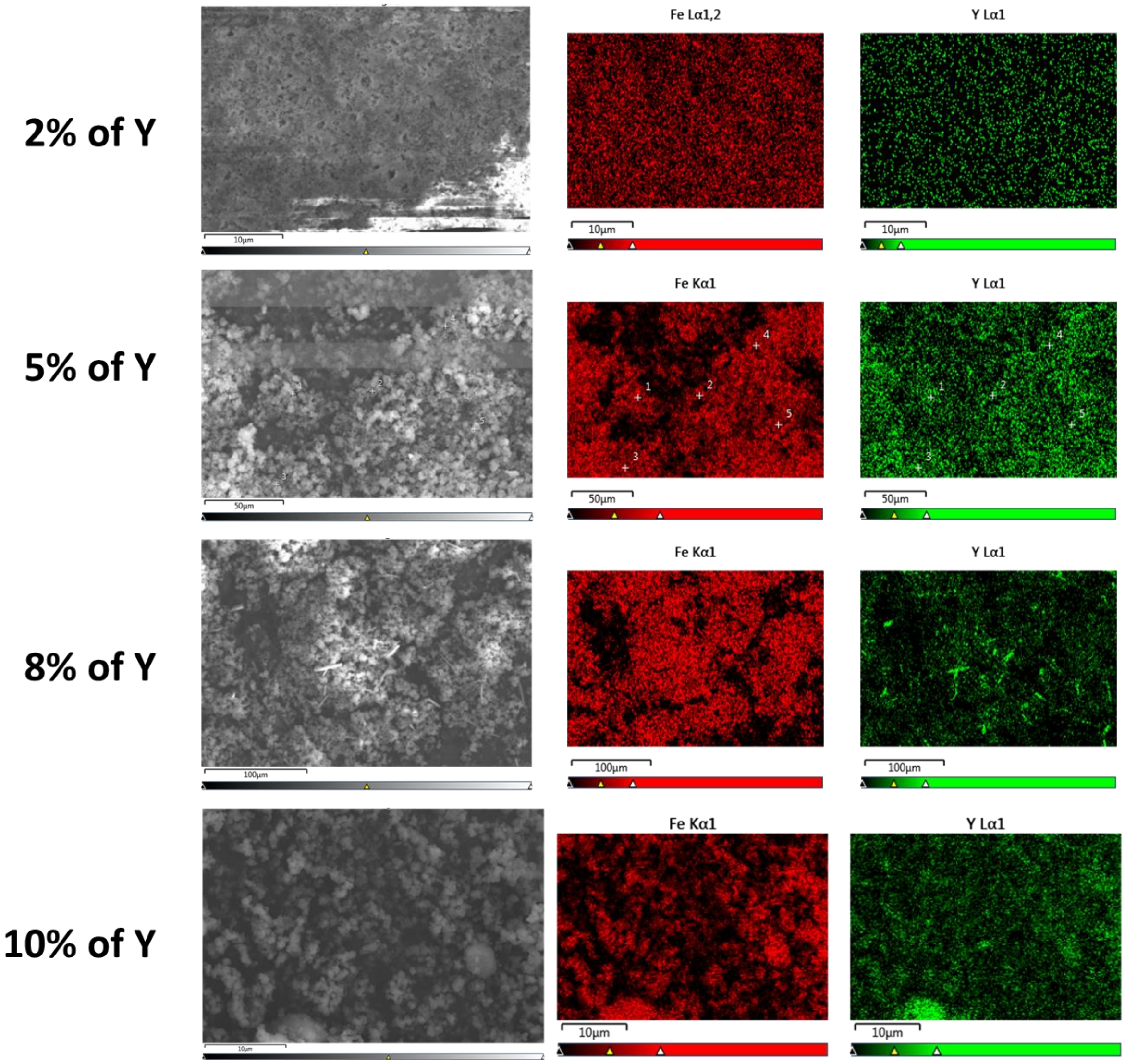


**Figure S5.** SEM EDX map of MIL-100(Fe,Y) particles with 2, 5, 8, and 10% of doped Y. The scale bar is 10 μm.

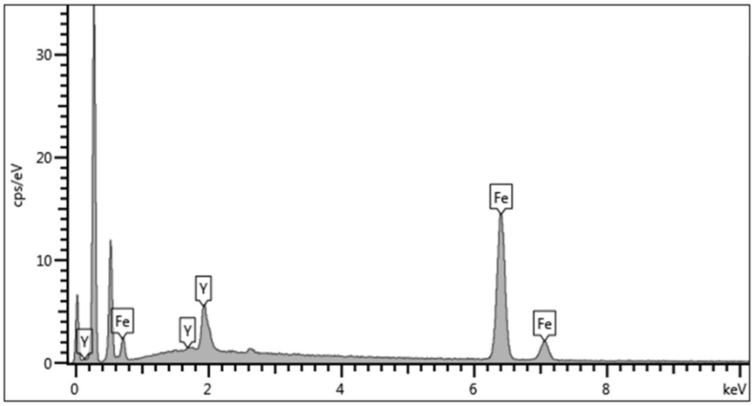


**Figure S6.** SEM-EDX spectrum of MIL-100(Fe, Y) with n(Y)/[n(Y)+n(Fe)] ratios at 10% of doped Y.

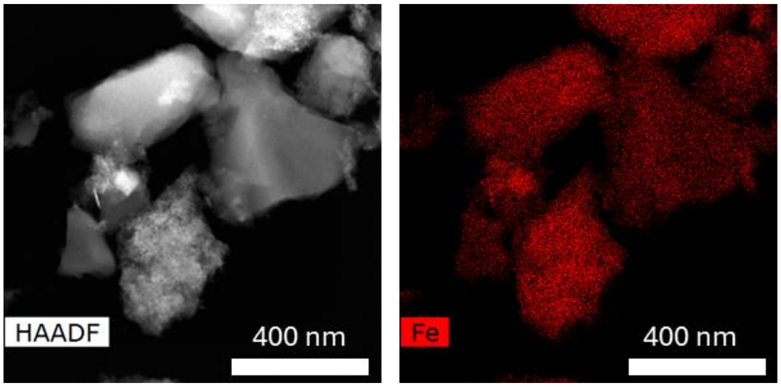


**Figure S7**. STEM-HAADF EDX map of MIL-100(Fe) particles.

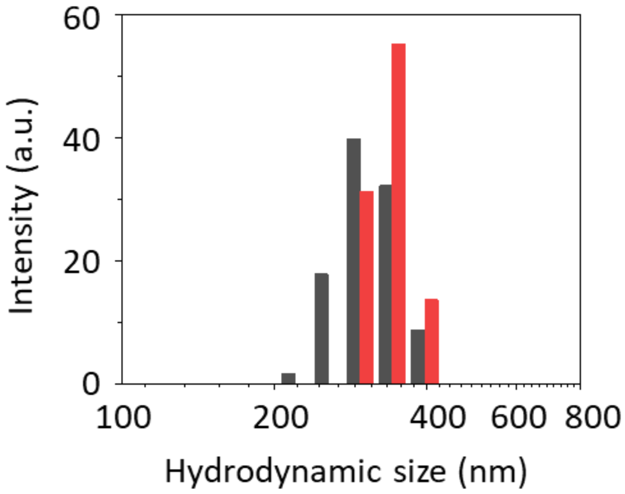


**Figure S8.** Hydrodynamic size of MIL-100(Fe) and MIL-100(Fe,Y) in PBS (pH=7.4).

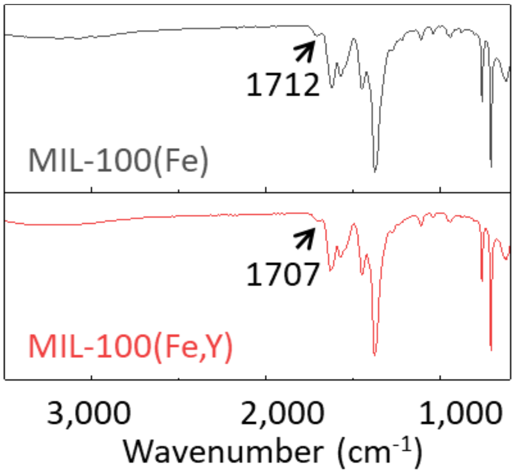


**Figure S9.** FT-IR spectra of MIL-100(Fe) and MIL-100(Fe,Y).

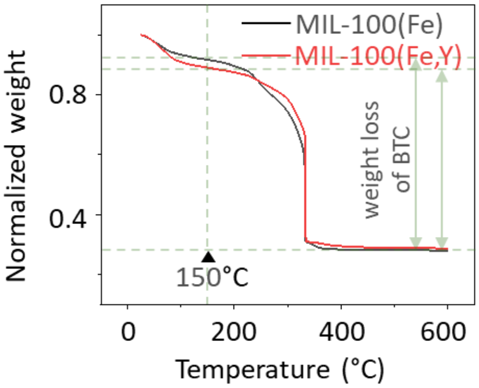


**Figure S10.** TGA curve of MIL-100(Fe) and MIL-100(Fe,Y), the mass of MIL-100(Fe) and MIL-100(Fe,Y) particles at 25 °C was normalized to "1".

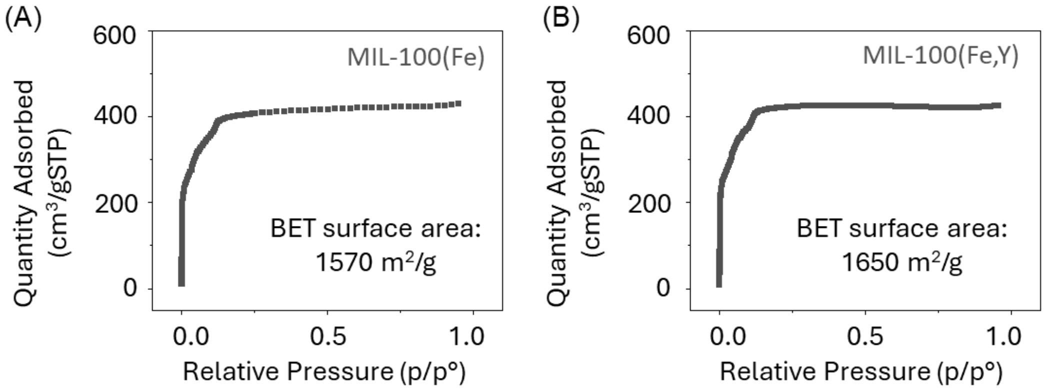


**Figure S11.** Nitrogen adsorption analysis of (A) MIL-100(Fe) and (B) MIL-100(Fe,Y).

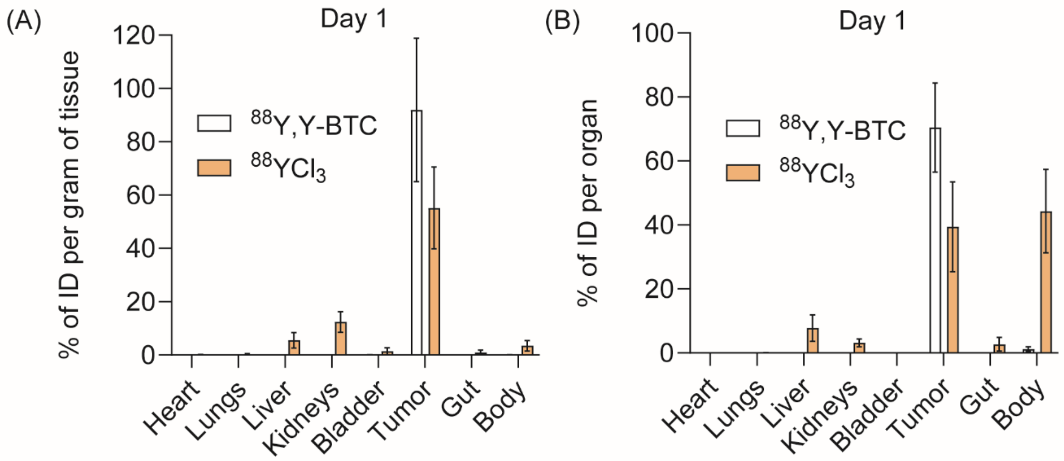


**Figure S12.** Yttrium-88 retention at tumor site after intratumoral injection of $^{88}$Y,Y-BTC-doped MOFs or $^{88}YCl_3$. (A) Radioactivity measurements in different tissues on day 1 after treatment with $^{88}$Y,Y-BTC particles compared to $^{88}YCl_3$ shown as the percentage of injected dose (ID) per gram of tissue and (B) per organ. The data are expressed as the mean ± SD (n = 3).

#### Supplementary Tables

**Table S1.** Comparison of Y molar ratios during synthesis and in resulting samples measured by SEM-EDX.

| n(Y)/[n(Y)+n(Fe)] (%) | |
|---|---|
| **during synthesis** | **measured by SEM-EDX ± error** |
| 2 | 2.39 ± 0.59 |
| 5 | 7.66 ± 1.22 |
| 8 | 10.50 ± 1.83 |
| 10 | 12.73 ± 1.58 |

**Table S2.** Intensity mean size and mean zeta potential of MIL-100(Fe) and MIL-100(Fe,Y) particles.

| Samples | Intensity mean size (nm) | PdI | Zeta potential (mV) |
|---|---|---|---|
| MIL-100(Fe) | 344.5 ± 27.3 | 0.7 | -23.0 ± 1.8 |
| MIL-100(Fe,Y) | 352.5 ± 91.5 | 0.7 | -24.3 ± 1.3 |

#### Supplementary Methods

*Cell culture*

The human adenocarcinoma alveolar epithelial A549 cell line (CCL-185), murine Lewis lung carcinoma (LLC-1) cell line (CRL-1642), and murine B16F1 melanoma cell line (CRL-6323) were obtained from ATCC. The A549 cells were cultured in DMEM/F12 medium supplemented with 10% FBS (v/v), 100 IU $mL^{-1}$ penicillin, 100 µg $mL^{-1}$ streptomycin, and 2 mM Gluta-max (Gibco) in a 95% humidified atmosphere containing 5% $CO_2$ at 37 °C. The LLC-1 cell line was cultured in DMEM, while the B16F1 cell line was cultured in DMEM/F12 medium supplemented with 10% FBS (v/v), 100 IU $mL^{-1}$ penicillin, and 100 µg $mL^{-1}$ streptomycin in a 95% humidified atmosphere containing 5% $CO_2$ at 37 °C.

*In vivo CT imaging*

To perform CT imaging, anesthetized mice with B16F1 tumors were injected with Y-BTC (100 mg $mL^{-1}$, 88 µL per 200 µL of tumor) in saline. Whole-body CT imaging was performed on a U-SPECT-II/CT scanner (MILabs, Utrecht, Netherlands) using a standard protocol with 45 kV and 500 µA. The CT imaging parameters were as follows: pixel size, 80 µm; field of view, 82 mm length × 41 mm diameter; rotation, 360°; step, 0.72°. The CT scans were collected and reconstructed using the manufacturer's software. PMOD 3.4 (PMOD Technologies Ltd., Switzerland) was used for 3D reconstruction and imaging.

*Dosimetry for treatments*

To calculate the absorbed radiation doses to tumors and cells from $^{90}$Y-containing formulations, we used the medical internal radiation dose (MIRD) method. In the MIRD schema, the absorbed dose $D_{\bar{r}_T}(\mathrm{t})$ to target tissue $\bar{r}_T$ over a dose integration period t was calculated as follows:

$$D_{\bar{r}_T}(\mathrm{t}) = \overline{\mathrm{A}}_{\bar{r}_S}(\mathrm{t}) \times \mathrm{S}(\bar{r}_T \leftarrow \bar{r}_S, \mathrm{t}),$$

where $\overline{\mathrm{A}}_{\bar{r}_S}(\mathrm{t})$ represents the time-integrated activity (i.e., the total number of nuclear transformations) in the source tissue $\bar{r}_S$, and $\mathrm{S}(\bar{r}_T \leftarrow \bar{r}_S, \mathrm{t})$ denotes the mean absorbed dose rate to the target region $\bar{r}_T$, at time t after administration per unit activity in the source region $\bar{r}_S$. For injected tumors, the source and target regions are the same. The S value is specifically determined for $^{90}$Y emissions based on the geometry and density of the tumor:

$$\mathrm{S}(\bar{r}_T \leftarrow \bar{r}_S, \mathrm{t}) = \frac{\sum_i E_i \cdot n_i \cdot \varphi(\bar{r}_T \leftarrow \bar{r}_S; E_i, t)}{m_{\bar{r}_T}(t)},$$

where $m_{\bar{r}_T}(t)$ represents the time-dependent mass of the target tissue, $n$ is a number of particles (or photons) with energy $E$ emitted per nuclear transformation in the radionuclide decay scheme, $\varphi(\bar{r}_T \leftarrow \bar{r}_S; E_i, t)$ is the absorbed fraction of energy $E$, which is emitted at time t from the source region $\bar{r}_S$ and is absorbed in the target volume $\bar{r}_T$. For our calculations, it was assumed that $\varphi(\bar{r}_T \leftarrow \bar{r}_S; E_i, t) = 1$.

*Histological analysis*

Tissues from the bone marrow, heart, lung, and liver were collected from two mice in the control group (on day 5) and two mice in the $^{90}$Y,Y-BTC microparticle-treated group (on day 11). The excised samples were placed in a 10% formalin solution for fixation. The tissues were then dehydrated, embedded in paraffin, cut into 5–6 μm-thick sections, and stained with hematoxylin and eosin. Analysis was performed using an Olympus CX41 microscope (Olympus Co., Tokyo, Japan), equipped with a UPlanApo 20×/NA 0.70 objective lens.
